# High plasticity of ribosomal DNA organization in budding yeast

**DOI:** 10.1101/2023.09.26.559284

**Authors:** Shuangying Jiang, Zelin Cai, Yun Wang, Cheng Zeng, Jiaying Zhang, Wenfei Yu, Chenghao Su, Shijun Zhao, Ying Chen, Yue Shen, Yingxin Ma, Yizhi Cai, Junbiao Dai

## Abstract

In eukaryotic genomes, ribosomal DNA (rDNA) generally resides as a highly repetitive and dynamic structure, making it difficult to study. Here, a synthetic rDNA array on chromosome III in budding yeast was constructed to serve as the sole source of rRNA. Utilizing the loxPsym site within each rDNA repeat and the Cre recombinase, we were able to reduce the copy number to as few as eight copies. Additionally, we constructed strains with two or three rDNA arrays, and found that the presence of multiple arrays did not affect the formation of a single nucleolus. Although alteration on the position and number of rDNA arrays did impact three-dimensional genome structure, the additional rDNA arrays had no deleterious influence on cell growth or transcriptomes. Together, this study sheds light on the high plasticity of rDNA organization and opens up opportunities for future rDNA engineering.

**Highlights:**

- A method was established for efficient construction of synthetic rDNA arrays in budding yeast
- The rDNA repeats in a haploid yeast can be reduced to as few as eight copies to support cell viability
- Yeast cells with two or three DNA arrays on distinct chromosomes form a single nucleolus.
- Dispersed rDNA arrays result in no deleterious influence on cell growth or transcriptomes.

## Introduction

Ribosomal DNA (rDNA) encodes genes for structural RNA components of the ribosome (Kobayashi, 2006). Due to the high cellular demand for ribosome biogenesis, most organisms have multiple copies of rDNA genes, and the rDNA region is highly transcribed to produce more than 70% of all cellular RNAs in a eukaryotic cell (Cerqueira and Lemos, 2019; Salim and Gerton, 2019). rDNA organization is largely conserved in most eukaryotes, with rDNA genes typically arranged in large stretches of tandem repeats (Nelson et al., 2019; Salim and Gerton, 2019). The repetitive rDNA arrays, termed nucleolar organizer regions (NORs), give origin to the nucleolus, a large dynamic membrane-less nuclear organelle (Cerqueira and Lemos, 2019). rDNA and the nucleolus are now understood to function as a coordinating hub for modulating diverse biological processes such as maintenance of epigenetic states, gene expression, cell proliferation, and aging (Cerqueira and Lemos, 2019; Hotz et al., 2022).

The tandem arrangement of repeats and high rates of transcription make rDNA one of the most unstable genomic regions and highly susceptible to recombination-mediated copy loss (Salim and Gerton, 2019). Despite this inherent instability, rDNA copy number is generally maintained in a certain range in each species, such as 350 copies in a human cell and 150 copies in a haploid yeast cell, illustrating the presence of mechanisms to recognize and regulate rDNA copy number (Kobayashi, 2006; Nelson et al., 2019). As a tractable model organism, budding yeast has been extensively used to explore the mechanisms to maintain stability of the rDNA locus (Nelson et al., 2019). Proteins such as Sir2 and Fob1 have been identified to be part of the pathway that sense loss of rDNA repeats and feed back into regulation of recombination and replication, thereby keeping rDNA copy number in a certain range (Iida and Kobayashi, 2019; Kobayashi and Sasaki, 2017; Nelson et al., 2019; Salim et al., 2017).

However, the hundreds or thousands of rDNA copies in eukaryotes are usually far in excess of the requirements for ribosome biogenesis and only a subset of rDNA repeats are transcribed even in actively growing cells of mammalians or yeast (Conconi et al., 1989; Iida and Kobayashi, 2019; Nelson et al., 2019; Salim and Gerton, 2019). By loss of native rDNA copies, viable yeast strains with low rDNA copy number in which all the rDNA copies were actively transcribed, were successfully constructed (French et al., 2003; Ide et al., 2010; Iida and Kobayashi, 2019; Takeuchi et al., 2003). Yeast strains with ∼20 copies of rDNA repeats showed normal expression levels of rRNA and ∼20% longer doubling time than wild type (>100 copies) (Ide et al., 2010; Takeuchi et al., 2003). Later, viable strains with 15 copies of rDNA repeats were constructed (Iida and Kobayashi, 2019). However, due to lack of sophisticate tools to engineer rDNA repeats, it remains untested whether rDNA copy number could be reduced even further.

Besides rDNA copy number, the number of rDNA arrays also varies among species. For example, in human, rDNA repeats are arranged into tandem arrays on five acrocentric chromosomes, and in mice on six (Cerqueira and Lemos, 2019). As for the single-cell organism *Saccharomyces cerevisiae*, the repeats are present in a single array on the right arm of chromosome XII (*chrXIIR*) (Salim and Gerton, 2019). From the rDNA database of plant and animals, most organisms have multiple rDNA arrays in their genome, with an average number of three to four loci per diploid genome (Garcia et al., 2017; Sochorova et al., 2018). The mechanisms governing rDNA loci number and the biological significance of multiple rDNA arrays remain unknown. Haploid yeast strains with two rDNA arrays have been constructed in a few studies (Hult et al., 2017; Lazar-Stefanita et al., 2023; Oakes et al., 2006). Building yeast models with three or more rDNA arrays, which more closely mimics the distribution patterns of higher organisms, has not been reported yet.

To probe the plasticity of rDNA organization, we engineered both rDNA copy number in a haploid strain with a single rDNA array and the number of rDNA arrays in a yeast cell. To efficiently construct rDNA arrays at target sites, we established a simple method that we termed <u>c</u>onstruction of a repetitive <u>a</u>rray by transformation (CAT, Fig.1A). Using CAT and a strain in which the native rDNA array on *chrXIIR* was eliminated, we successfully built a synthetic rDNA array on *chrIIIR* acting as the sole source of rRNA. A loxPsym site was inserted in each repeat. Through induction of rDNA array rearrangement mediated by Cre recombinase, we found that copy number of rDNA repeats could be reduced to only eight copies. In addition, we constructed strains with two or three rDNA arrays at diverse genomic loci. Although obvious changes of chromosome conformation were identified in the three-dimensional (3D) genome structure by Hi-C, the existence of additional rDNA arrays did not disrupt cell growth, global gene transcription or nucleolar structure. Our results indicate that rDNA in budding yeast tolerates radical changes in organization and provide tools for rDNA engineering in synthetic genomes.

**Figure 1.**
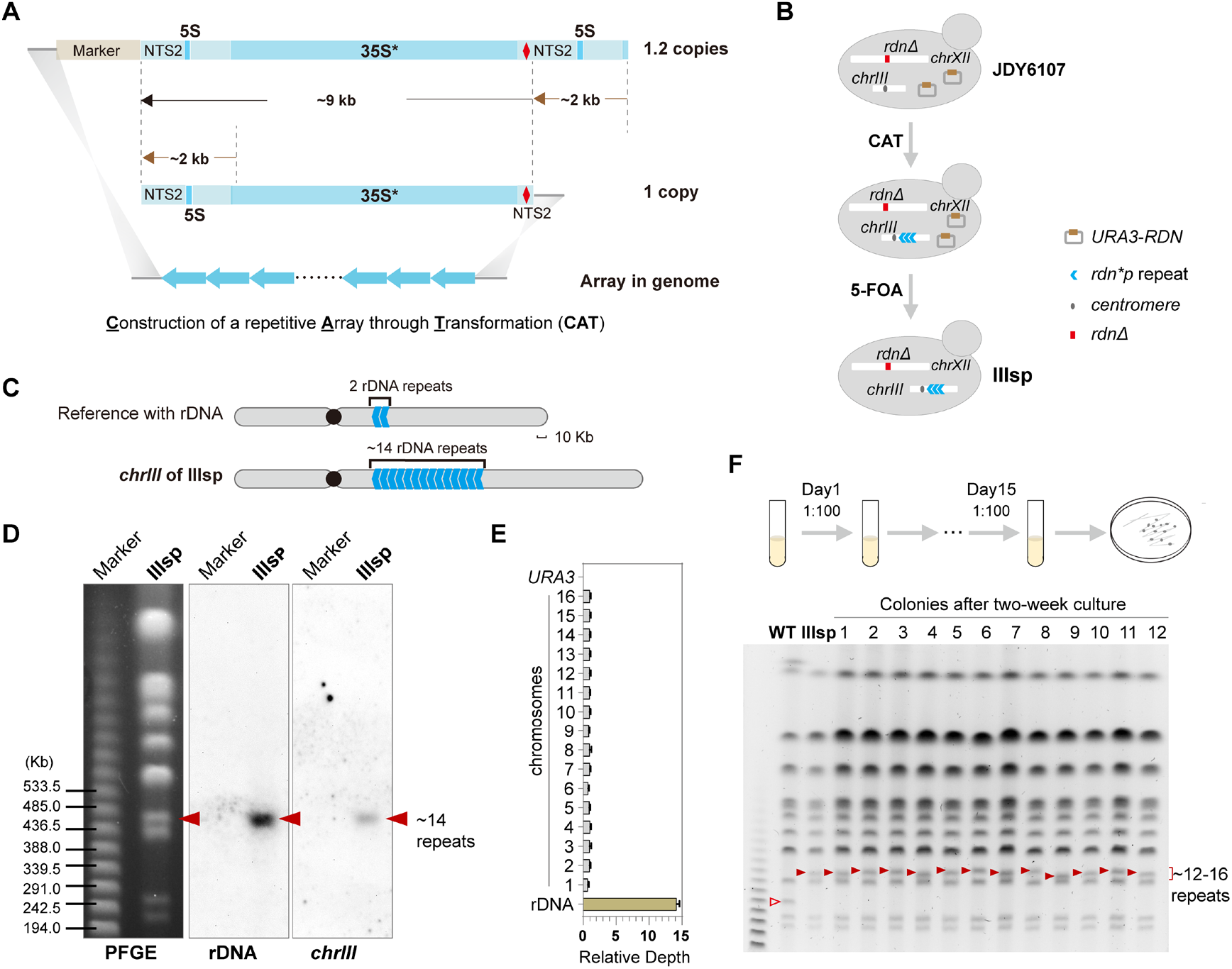
Construction of a short synthetic rDNA array on *chrIII*. **(A)** <u>C</u>onstruction of a repetitive <u>A</u>rray through <u>T</u>ransformation (CAT).*: a *hyg1* mutation in 18S. Red diamonds: loxPsym cassette inserted at position -206. For the plasmid containing 1.2 copies rDNA repeats, a ∼2 kb rDNA fragment was placed downstream of a complete repeat (∼9 kb). (B) The design to construct the strain IIIsp. IIIsp designates the strain containing a single synthetic <u>s</u>hort rDNA array with lox<u>P</u>sym sites. *RDN*: the rDNA repeats with wild type sequences. *rdn*p* designates the rDNA repeat containing both a loxPsym site (p) and a *hyg1* mutation (*). (C) Diagram of *chrIII* in IIIsp based on the assembled contigs. The synthetic rDNA array had been correctly inserted in the strain IIIsp. (D) PFGE and southern blotting analysis of the integration of rDNA repeats on *chrIII*. Marker: NEB lambda PFG ladder. Probes for rDNA and *chrIII* were used for southern blot. (E) The rDNA copy number of the IIIsp strain. The means of the actual Nanopore sequencing depth of a 1 kb-sized bin for all chromosomes in each strain were normalized as 1. The error bar represents the standard deviation (SD). (F) The stability of the synthetic short rDNA array. IIIsp was cultured in YPD medium for two weeks and then the streaked colonies were analyzed with PFGE. Red arrowheads: the bands of *chrIII*.

## Results

### Construction of a repetitive array by transformation

To engineer the rDNA organization, we need to have the capacity to efficiently build rDNA arrays with designed sequences at chosen target sites. To this end, we developed a method called <u>c</u>onstruction of a repetitive <u>a</u>rray by transformation (CAT, Figure 1A). It is based on the transformation of two linear DNA fragments. One fragment contains a single rDNA repeat (about 9 Kb), flanked by the sequence homologous to downstream of the targeted locus on the right side. The second fragment has 1.2 copies of rDNA repeats, a marker gene, and the sequence homologous to upstream of the targeted locus on the left side. Through homologous recombination, an rDNA array with a random number of repeats can be constructed upon transformation.

As a proof-of-concept, we built an additional rDNA array in the wild type strain BY4742 on the right arm of chromosome XV (*chrXVR)* at a site previously chosen for introduction of a synthetic rDNA array (Zhang et al., 2017). Two strains subjected to CAT with the targeted integration (strain a and b) were randomly selected and analyzed by southern blotting using probes specific to either rDNA or *chrXV*. About 15 rDNA repeats were integrated in strain b and fewer in a (Figure S1A). Given that 15 rDNA repeats have been reported to be sufficient for strain viability (Iida and Kobayashi, 2019), our results underscore the potential of CAT to build short functional rDNA arrays.

### A single short synthetic rDNA array in which each repeat contains a loxPsym site can support cell viability

In previous studies, several groups successfully reduced the rDNA copy number from typical ∼150 copies to 15-20 copies, the number which was thought to be close to the minimum number of rDNA copies to support cell growth (Ide et al., 2010; Iida and Kobayashi, 2019; Takeuchi et al., 2003). As a unique system implanted in the synthetic yeast genome, the SCRaMbLE (synthetic chromosome rearrangement and modification by loxP-mediated evolution) system has been demonstrated to be effective for chromosome minimization (Luo et al., 2021). To investigate whether the rDNA copy number could be further reduced, we wanted to introduce the Cre-loxPsym system into rDNA repeats. The high-copy plasmid *LEU2-RDN* harboring a single wild-type rDNA repeat was modified by introducing a loxPsym site in the nontranscribed spacer 2 (NTS2) region (Figure S1B). The resulting plasmid with an engineered rDNA repeat was named *LEU2-rdnp*. *LEU2-rdnp* or *LEU2-RDN* was transformed into the strain JDY6106, whose entire native rDNA array on *chrXIIR* has been deleted (*rdnΔ*) and a high-copy plasmid containing a *RDN* repeat (*URA3-RDN*) was the sole source of rRNA. Plasmid shuffling by treatment with 5-FOA could remove *URA3-RDN,* and *LEU2* marked plasmids containing functional rDNA repeats will support cell growth. Like the *RDN* repeats, the *rdnp* repeats supported cell growth, indicating insertion of the loxPsym site did not disrupt the functions of rDNA repeats (Figure S1B).

To make a synthetic rDNA array with designed sequences the sole source of rRNA, JDY6106 was again used as the target strain (Figure 1B). To stabilize the integrated rDNA array, the gene *FOB1*, which is required for rDNA expansion (Johzuka et al., 2006; Kobayashi et al., 1998; Kobayashi et al., 2001), was also deleted in JDY6106, generating JDY6107. The recessive *hyg-1* mutation in 18S rRNA gene, which was used to induce rDNA expansion through hygromycin B selection (Chernoff et al., 1994; Zhang et al., 2017), was introduced into rDNA repeats in addition to loxPsym sites (*rdn*p*, Figure 1A and 1B). A locus on *chrIIIR* previously used for integration of a synthetic rDNA array (Zhang et al., 2017) was chosen as the insertion site. After applying CAT to JDY6107 with the two linearized fragments containing *rdn*p* repeats (Figure 1A and 1B), cells were grown in media containing 5-FOA to identify clones in which the *URA3-RDN* plasmids had been lost. Six clones were randomly selected and analyzed with southern blotting. We found multiple rDNA repeats were successfully incorporated on *chrIII* (>11 copies) in all strains and there was no rDNA at other chromosomal locations (Figure S1C).

One of the six strains was randomly chosen and named as IIIsp. Results of whole genome sequencing using Nanopore long-read sequencing and southern blotting confirmed the deletion of rDNA array from *chrXII* and the insertion of a synthetic array on *chrIII* (Figure 1C-D, S1D-1E). Judging by the length of the new *chrIII* (Figure 1D), approximately 14 copies of rDNA repeats were inserted. Read depth is another way reported to estimate rDNA copy number (Gibbons et al., 2014; Tubbs et al., 2018). Similarly, we divided the average depth of the rDNA repeats by that of all chromosomes without rDNA to calculate the average copy number of rDNA repeats per haploid genome. This method also suggested an average rDNA copy number of IIIsp of about 14 (Figure 1E). These results confirmed that we successfully build a short synthetic rDNA array (∼14 copies) to replace the native long array on *chrXIIR*, yielding viable cells.

To examine the stability of the short synthetic rDNA array, IIIsp was inoculated in rich medium and subcultured daily for two weeks. Twelve colonies were randomly picked and analyzed (Figure 1F, S1G-1H). All isolates were still haploid ploidy (Figure S1F). No obvious growth difference was detected between IIIsp and these isolates, irrespective of whether they were cultured in rich medium at 30°C or at high temperature (Figure S1G). Compared to IIIsp, the *chrIII* bands of these isolates showed a slight up- or down-shift (Figure 1F). The variation of rDNA copy number in these strains is approximately from 12 to 16 copies. Together, the synthetic rDNA array is relatively stable in cells and the spontaneous reduction of rDNA copy number from 14 to 12 suggests the minimal rDNA copy number for cell viability may be lower.

### IIIsp exhibits normal nucleolus structure, cell morphology and cell size

In budding yeast, the nucleolus formed by the rDNA array can be identified as an approximately crescent-shaped structure positioned adjacent to the nuclear envelope (Hult et al., 2017). To visualize the nucleolus, Nop10-GFP and Nic96-mCherry were used to label the nucleolus and nuclear envelop, respectively, as reported previously (Zhang et al., 2017). Strains labeled with fluorescent proteins were named by adding an “F” after the original names. It has been reported that dispersed nucleolus signals were observed in *rdnΔ* cells bearing rDNA on a high-copy plasmid (Oakes et al., 1998; Zhang et al., 2017). Consistently, the strain JDY6107-F showed disruption of the well-organized nucleolar structure (Figure 2A). As for IIIsp-F, cells kept the short length of the ectopic rDNA array (∼13 copies) and formed a nucleolar structure indistinguishable from those of BY4742-F and *fob1Δ*-F (Figure 2A and S2A).

**Figure 2.**
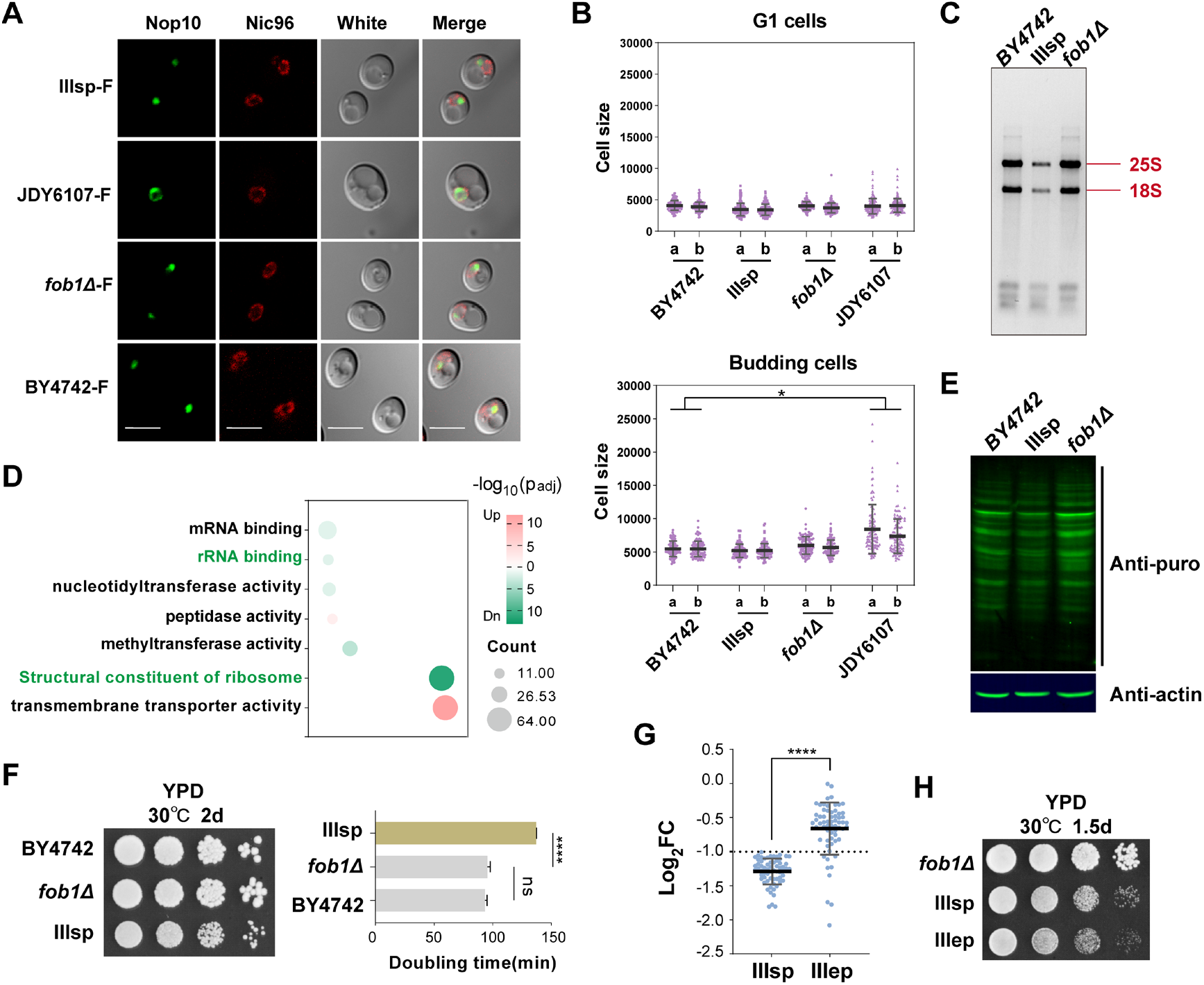
IIIsp exhibited normal nucleolus structure and attenuated global translation activity. (A) Microscopy pictures of the strains tagged with Nop10-GFP and Nic96-mCherry. The bar means 5μm. (B) Cell size quantification of the strains. The dot plot showed the distribution of cell sizes of G1 cells (a single cell without bud) and budding cells. Cells from two clones of each strain were counted. More than 100 cells for each clone at different states were counted. Cell sizes were measured by ImageJ. The mean and SD were shown. Nested one-way ANOVA was used to compare different groups. *represents P<0.05. (C) The total RNA extracted from the same number of cells was compared. (D) Molecular function enrichment analysis of the differentially expressed genes in IIIsp. Differential expression in this study was defined as |log_2_ (Fold Change)| > 1 and -log_10_ (Adjusted p-value) > 3. Sorting by -log_10_ (Adjusted p-value), the seven terms with adjusted p-value <0.05, including five down-regulated terms (green) and two up-regulated terms (pink), were represented. The bubble size represents gene count. (E) Global translation activity was measured by puromycin incorporation. β-actin was shown as the loading control. (F) The growth of different strains in rich media. Four technical replicates were conducted to calculate doubling time of the strains. Error bars represent SDs. Significance of differences was tested by two-sided t-tests. ****indicates P<0.0001. ns indicates no significance. (G) The expression of the 64 ribosomal protein genes in different strains. BY4742 was used as the control. Significance of differences was tested by two-sided t-tests. ****indicates P<0.0001. (H) Increase of rDNA copy number did not rescue the growth defects from the synthetic rDNA array.

Since we observed many enlarged budding cells in JDY6107-F, we also examined the effects of the short synthetic rDNA array in IIIsp on cell morphology and cell size. IIIsp showed normal cell morphology and did not differ in cell size to the control strains BY4742 and *fob1Δ*, which contained the native long rDNA array on *chrXIIR,* while JDY6107 showed a significant increase in cell size of budding cells (Figure 2B and S2B). Our above results indicate that decrease of rDNA copy number and translocation of a single rDNA array to another chromosome did not affect nucleolus structure, cell morphology or cell size.

### IIIsp shows reduced global translation activity and growth defects

Given that rRNAs are structural components of ribosomes, the short synthetic rDNA array could lead to defects in ribosome biogenesis or translation. To visualize the changes in the amount of rRNAs, total cellular RNA from the same number of cells was separated on agarose (Figure 2C). Compared to BY4742 and *fob1Δ*, IIIsp exhibited an obvious reduction of 25S rRNA and 18S rRNA level (Figure 2C). Besides rRNAs, ribosomal proteins are constituent components of ribosomes. To test whether expression levels of ribosomal proteins were affected, the transcriptomes of IIIsp and *fob1Δ* were compared to that of BY4742. For IIIsp, differentially expressed genes (DEGs) were enriched in the molecular functions involving structural constituents of the ribosome and rRNA binding. 64 ribosomal protein genes (∼50% of all ribosomal protein genes) were significantly downregulated (Figure 2D, Table S1). In addition, the two enzymatic subunits of RNA polymerase I, responsible for the synthesis of the precursor 35S rRNA (Engel et al., 2013; Salim and Gerton, 2019), namely *RPA135* and *RPA190*, were also significantly repressed in IIIsp (Table S1). Together, these results provide evidence for the defects of ribosome biogenesis in IIIsp. Since only 15 DEGs were identified in *fob1Δ* with none of the ribosomal protein genes being affected (Table S1), the defects in IIIsp evidently result from the short synthetic rDNA array. Global translation activity in IIIsp was analyzed with the puromycin incorporation assay as described previously (Hu et al., 2022). Puromycin incorporation signals of IIIsp were weaker than BY4742 and *fob1Δ*, indicating defects of IIIsp in global protein synthesis (Figure 2E).

Since ribosome biogenesis and protein synthesis drive cell growth (Lempiainen and Shore, 2009; Shore and Albert, 2022), we evaluated the growth of IIIsp. It has been reported that relative to strains carrying a typical copy number (>100 copies), low copy strains (15-20 copies on *chrXII*) grew more slowly (∼20% longer doubling time in rich medium) (Ide et al., 2010; Takeuchi et al., 2003) and ∼40 copies of wild type rDNA repeats were sufficient to maintain normal growth rate (French et al., 2003). We found that IIIsp also grew more slowly on YPD than BY4742 and *fob1Δ*: the doubling time of IIIsp showed a ∼50% increase compared to BY4742 and *fob1Δ* (Figure 2F).

One notable feature in IIIsp is the low rDNA copy number. To analyze the influence of rDNA copy number on strain phenotypes, we expanded the synthetic rDNA array in IIIsp to about 40 copies through the transient expression of *FOB1* and named the resulting strain IIIep. The strains with an expanded array on *chrIII*, IIIep-F, also showed normal nucleolar structure (Figure S2C-D). With increased rDNA copy number, the expression of ∼86% (55/64) down-regulated ribosomal protein genes in IIIsp was recovered to levels that did not significantly differ from BY4742 (Figure 2G, Table S1). However, about three quarters of DEGs in IIIsp (603/880) were also identified in IIIep (Figure S2E, Table S1). Using the method called spatial analysis for functional enrichment (SAFE) (Baryshnikova, 2016), these 603 DEGs were found to be enriched in rRNA assembly and maturation, DNA replication and repair, cell cycle and protein glycosylation (Figure S2F). In addition, IIIep still showed a growth defect on YPD comparable to that of IIIsp (Figure 2H). These results reveal that the low rDNA copy number may not be the main reason for the defects in IIIsp.

### rDNA copy number in haploid strains can be reduced to only eight copies

In the synthetic rDNA array in IIIsp, each repeat contains a loxPsym site, allowing to alter rDNA copy number via Cre-mediated recombination. A plasmid with Cre driven by a GAL inducible promoter was introduced into IIIsp (Figure 3A), leading to growth variations among resulting clones. We randomly picked six clones which grow either faster or slower than IIIsp on YPD for analysis. We found all clones with faster growth showed diploidy, drawing our attention to the ones with slower growth. Interestingly, we discovered that these slow-growing clones exhibited divergent patterns of *chrIII* in PFGE and southern blotting, including up or down-shifted, or even multiple bands, suggesting complicated rearrangement events have happened (Figure 3A). The two strains with an extremely short *chrIII*, S1 and S5, appeared to be diploid strains (Figure 3B). Two haploid strains with a shortened *chrIII*, S2 and S3, were subjected to Nanopore sequencing to assess the rDNA copy number, which revealed that the average rDNA copy number was about 7 for S2 and 9 for S3, respectively (Figure 3C). Using the long reads detected, we were able to assemble the sequence structure of *chrIII* in both S2 and S3 (Figure S3A). The reads that go across the whole arrays were not identified. To further define the rDNA copy number in these strains, the genomic DNA from a single colony of S2 was subjected to XmaI restriction digestion, followed by PFGE and southern blotting. Our control strain, IIIsp, yield an rDNA copy number of 14 (Figure 3D and S3B), consistent with results in Figure 1. In contrast, the chosen S2 colony contained 8 copies of rDNA repeats (Figure 3D).

**Figure 3.**
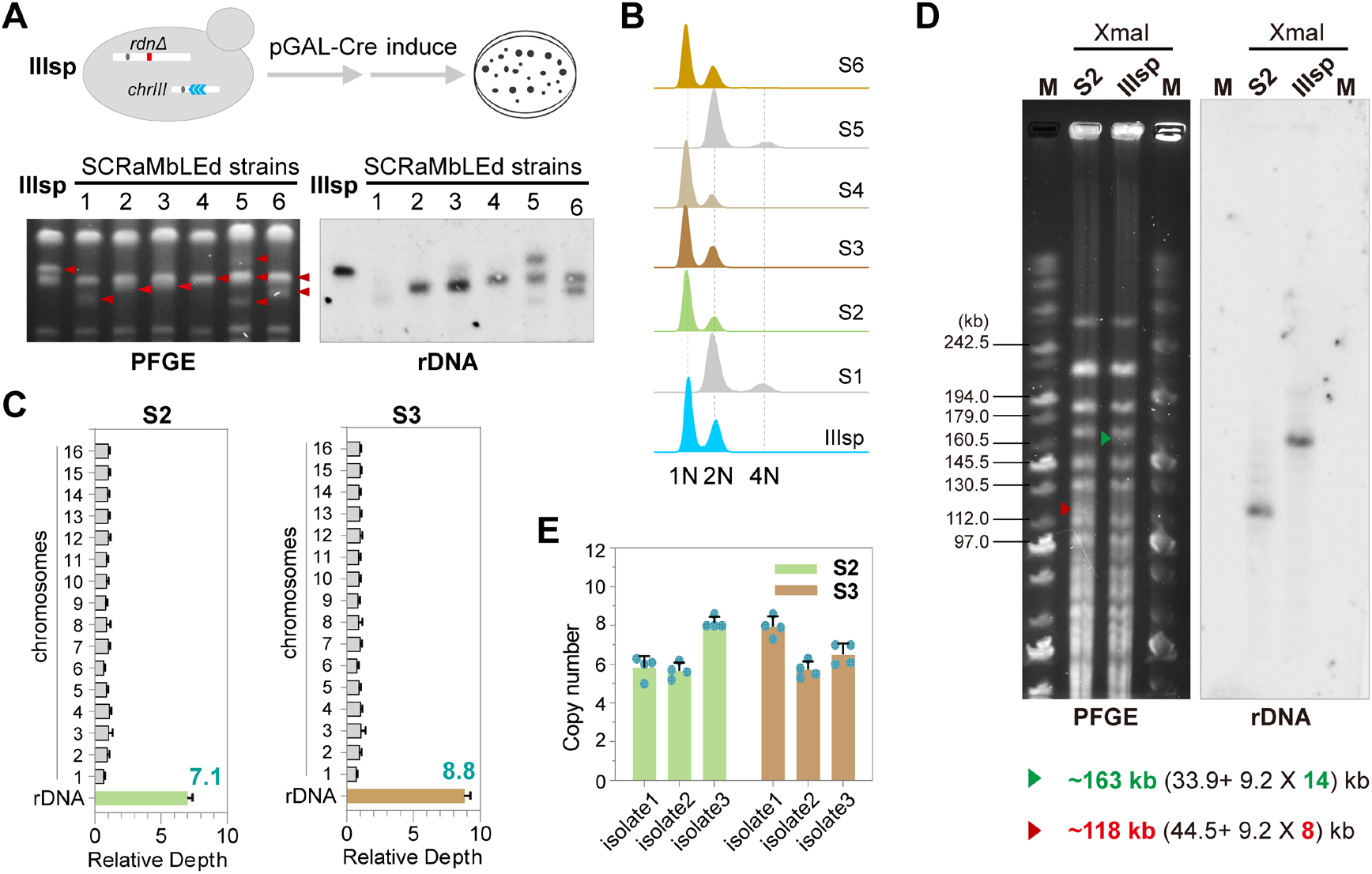
Generation of viable strains with approximately eight rDNA copies using SCRaMbLE. (A) PFGE and southern blotting analysis of the rDNA repeats on *chrIII* in the six strains generated by SCRaMbLE. IIIsp was used as the control. Specific probes for rDNA were used for southern blotting. Red arrowheads: the bands of *chrIII*. (B) Ploidy analysis of the generated strains using asynchronous log-phase cells. The haploid control strain IIIsp was used. S1-S6: represent the SCRaMbLEd strains in (A). (C) The rDNA copy number of the SCRaMbLEd strains was determined by relative read depth, as in Figure 1. (D) PFGE and southern blotting analysis of the rDNA copy number in a single colony of S2. The control strain IIIsp and specific probes for rDNA were used for southern blotting. NEB MidRange PFG Marker (M) was used. Red and green arrowheads: The bands of the rDNA fragment in S2 and IIIsp. (E) The rDNA copy number of the SCRaMbLEd strains evaluated by ddPCR. Three isolates from each strain were analyzed with the errors bar representing SD of four technical repeats.

To obtain an accurate number of rDNA repeats, we turned to droplet digital PCR (ddPCR), which eliminates the need for standard curves and provides copy number of a gene target on an absolute scale with high precision (Salim et al., 2017). The rDNA copy number per haploid genome was defined as the ratio of the absolute number of 25S gene to that of *TUB1* in the same sample. Given the rDNA copy number was more stable in stationary-phase (Figure S3C), we limited all subsequent analysis in cells at this stage. We analyzed three random isolates of both S2 and S3 (Figure S3D) and found that each strain contained 6–8 rDNA repeats (Figure 3E). Collectively, these results demonstrated that rDNA copy number of haploid strains can be reduced to as few as approximately eight copies.

### The existence of additional rDNA arrays does not disrupt normal cell growth

Unlike budding yeast, higher organisms usually contain multiple rDNA arrays in different chromosomes (Garcia et al., 2017; Sochorova et al., 2018). However, the evolutionary implications of possessing multiple rDNA arrays remain obscure. To emulate this multi-array configuration, we built yeast strains with two or three rDNA arrays located on separate chromosomes (Figure 4A). Given the high sequence identity amongst multiple rDNA arrays, inter-chromosomal recombination may occur. To effectively track these recombination events between rDNA arrays, we used the loxPsym cassette within *rdn*p* repeats as a distinctive tag. Specifically, we generated an expanded rDNA array with *rdn*p* repeats on *chrIIIR* in the strain where the native rDNA array on *chrXIIR* had been deleted (strain JDY6117, Figure 4A and S4A). Then, strain b, generated by CAT, was employed as one of the parental strains and renamed as XV-XII, denoting the positions of rDNA arrays (Figure S1A). Next, the two aforementioned strains, JDY6117 and XV-XII were crossed to produce the diploid strain Dip3 which now harbors three rDNA arrays (Figure 4A). Finally, Dip3 was sporulated, generating haploid strains containing single, two, or three rDNA arrays (Figure 4A and S4B). These new strains were designated as III, XV, III-XV, III-XII and III-XV-XII, following the naming convention established for XV-XII. Additionally, the wild-type strain BY4742 was named as XII.

**Figure 4.**
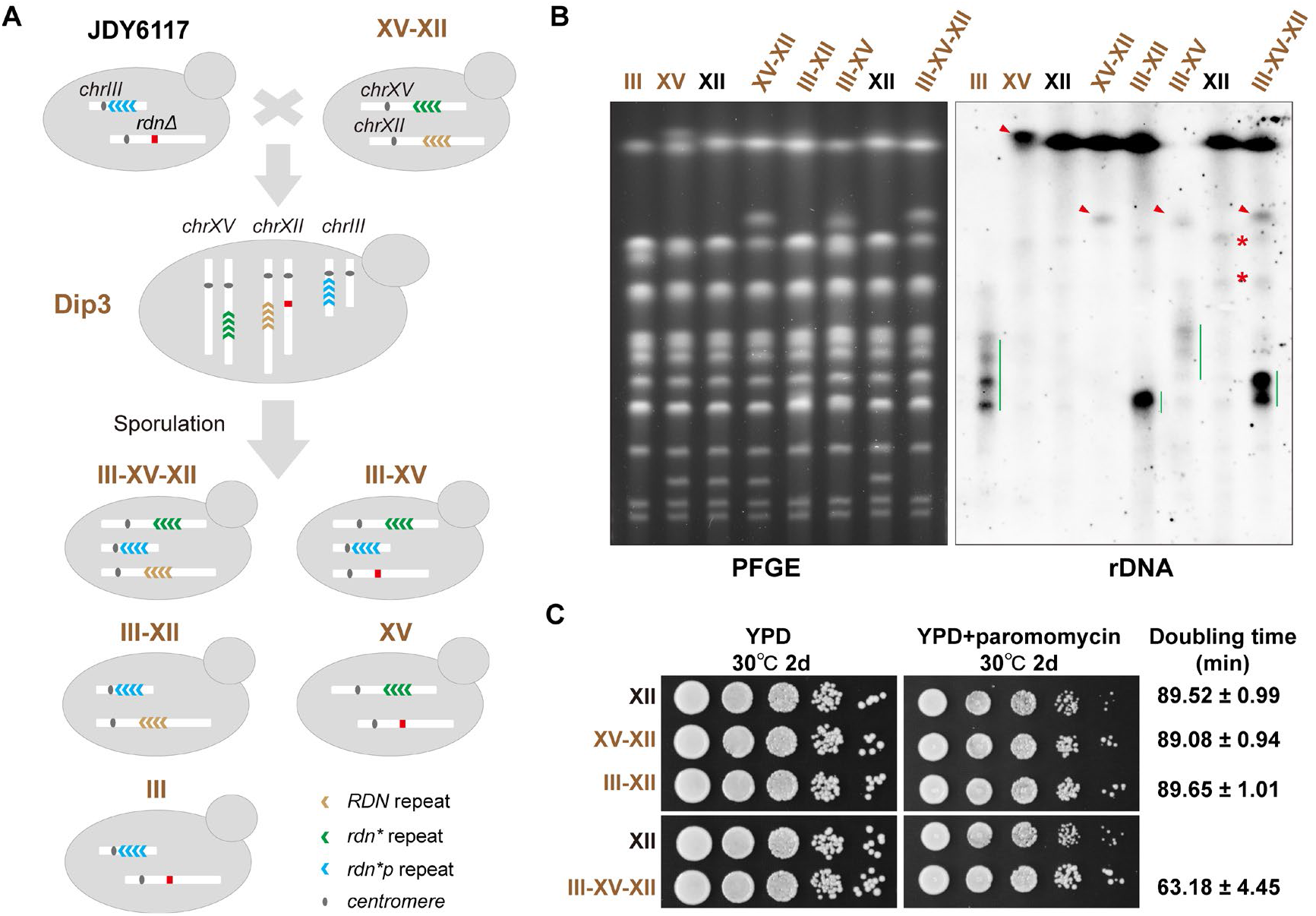
The existence of additional rDNA arrays did not impair cell fitness. (A) Generation of strains with different numbers of rDNA arrays. Dip3: the diploid strain with triple rDNA arrays on different chromosomes. *Rdn\** repeat: the rDNA repeat containing a *hyg1* mutation in 18S(*). (B) PFGE and southern blotting analysis of the strains with different numbers of rDNA arrays. Specific probes for rDNA were used for southern blotting. Red arrowheads: the bands of *chrXV* with rDNA repeats. Green lines: the signals of *chrIII* with rDNA repeats. Red stars: the unspecific signals. XII: BY4742. (C) Spot assays of different strains on rich media. Three technical replicates were cultured in YPD to calculate doubling time of the strains. The average doubling time ±SD is shown in the right.

The positioning of rDNA arrays in above strains has been verified through long-read sequencing, revealing no duplication of a particular chromosome within these strains (Figure S4C). The reads containing rDNA repeats can be categorized into two types, namely the rDNA-only reads and those containing both rDNA repeats and flanking genomic regions (Figure S5). It is worth noting that our strategy for strain construction ensured that the rDNA repeats with loxPsym sites solely originated from the array on *chrIIIR*. Consistently, no loxPsym sites were identified in the rDNA repeats of the parental strain XV-XII that had no access to *rdn*p* repeats. In the cases of strains carrying a single rDNA array, each rDNA repeat in III featured a loxPsym site while no loxPsym sites were identified within the rDNA repeats of XV. We have thoroughly analyzed the reads that included rDNA repeats and the neighboring genomic regions from III-XV, III-XII and III-XV-XII, and were unable to identify any re-location events of the loxPsym sites to *chrXII* or *chrXV*. Our analysis of hundreds of rDNA-only reads demonstrated that nearly all rDNA repeats present in a certain read were homogenous (Figure S5). These results collectively suggest the rarity of inter-array recombination events during the construction of all five strains through mating and meiosis.

To analyze the length of rDNA arrays on different chromosomes, we employed PFGE and southern blotting techniques (Figure 4B). The diverse rDNA signals observed in the parental strain JDY6117 were in agreement with the varying lengths of *chrIII* harboring rDNA repeats in III, III-XII, III-XV and III-XV-XII (Figure 4B and S4A). The length of rDNA array on *chrXVR* in III-XV and III-XV-XII was akin to that in XV-XII. Conversely, the rDNA array on *chrXVR* within XV was markedly expanded, aligning with the estimated rDNA copy number (∼55 copies) determined in Figure S4C.

Using above strains, we evaluated the effects of an additional array (XV-XII or III-XII) or two additional arrays (III-XV-XII) on cell growth. The rDNA array located on *chrXVR* in XV, acting as the exclusive rRNA source within the genome, supported cell growth as efficiently as the native rDNA array on *chrXIIR* in BY4742, while the *rdn*p* array on *chrIIIR* in III led to a slow-growth phenotype reminiscent of IIIep (Figure S6A). It is notable that regardless of the position and sequences of the additional rDNA array, both XV-XII and III-XII exhibited comparable colony size and doubling time (∼90min) to BY4742 (XII) in rich medium (Figure 4C). Strikingly, when compared to BY4742, the strain carrying two additional rDNA arrays, III-XV-XII, displayed similar colony size on YPD plates, but a much shorter doubling time (63.18 ± 4.45min, Figure 4C and S6B). In addition, under the treatment of paromomycin, XV showed a greater colony size than BY4742, suggesting that the *hyg-1* mutation in rDNA repeats augmented resistance to the translation error-inducing drug (Figure S6A). Consistently, XV-XII, III-XII and III-XV-XII, the strains harboring the *hyg-1* mutation, all showed increased resistance to paromomycin (Figure 4C). Together, the presence of additional rDNA arrays does not disrupt normal cell growth, but instead potentially confers growth rate benefits.

### Multiple rDNA arrays in yeast strains form a single nucleolus

In higher eukaryotic cells, the number of nucleoli is usually smaller than the number of rDNA arrays (NORs), and several NORs are often found within a single nucleolus, e.g., 10 NORs vs 2-5 nucleoli in Hela cells (Cerqueira and Lemos, 2019; Lafontaine et al., 2021). Given the presence of multiple rDNA arrays, we investigated whether the yeast strains would form multiple-nucleolar structures. To avoid the influence of cell division on the number of nucleoli, only G1 cells were counted. Using the same fluorescent proteins in Figure 2A, we found that nearly all cells (>97%) of the haploid strains with the number of rDNA arrays ranging from 1–3 displayed only a single, well-structured nucleolus (Figure 5A-B). In diploid strains, Dip3-F cells showed a 96.6 ± 0.3 % frequency of a single nucleolus structure, comparable to the diploid control BY4743-F (97.2 ± 1.1 %, Figure S7A-B). These results illustrate that multiple rDNA arrays in yeast form a single nucleolus.

**Figure 5.**
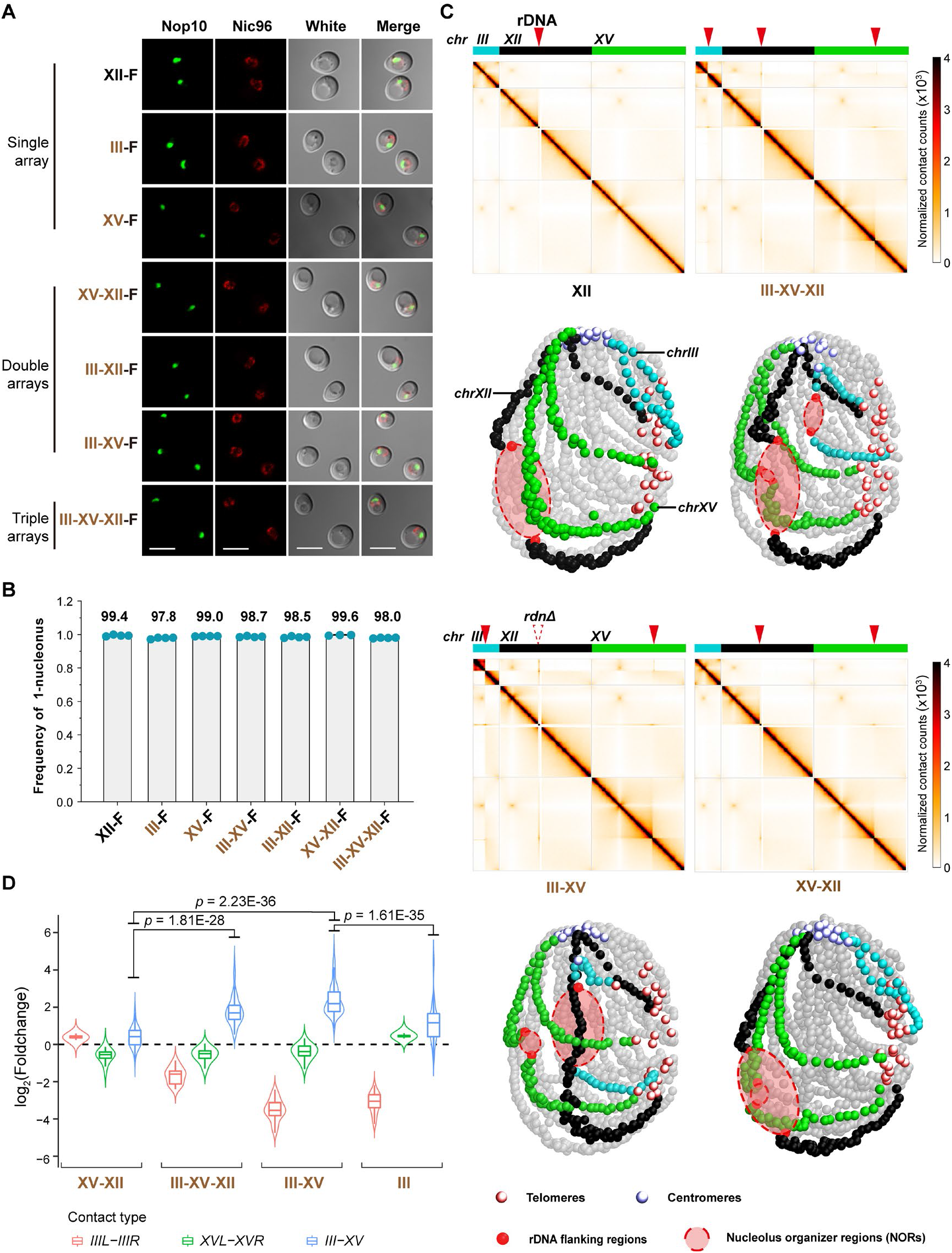
Yeast strains with multiple rDNA arrays exhibit a single nucleolus. (A) Microscopy pictures of the strains tagged with Nop10-GFP and Nic96-mCherry. The bar corresponds to 5μm. (B) The frequency of a single nucleolus in cells with different number of rDNA arrays. More than 200 un-budding cells were calculated for each clone. Three or four independent clones were analyzed for each strain (blue dots) and the average frequency were listed above. (C) Effects of rDNA arrays on genome structure. Upper panels show the Hi-C contact maps of the three chromosomes (*chrIII*-blue, *chrXII*-black, *chrXV*-green) in corresponding strains. The solid red triangle points to the position of the rDNA arrays and the dotted triangle points to the position of deleted native rDNA array on *chrXII*. The lower panels show the 3D representations of corresponding contact maps. The telomeres, centromeres, rDNA flanking regions (10kb) and nucleolus organizer regions (NORs) were represented on each structure. (D) Comparison of contact frequency between different rDNA flanking regions (50kb) in different strains to that of BY4742. *IIIL-IIIR* (red) and *XVL-XVR* (green): the interactions between upstream and downstream sequences of the rDNA insertion site on *chrIII* or *chrXV*. *III-XV* (blue): the interactions between the flanking regions on *chrIII* and those on *chrXV*. Significance of differences was tested by two-sided t-test, p-value was listed above.

Given the observation that the haploid strains with two or three rDNA arrays displayed single nucleolus, it is reasonable to hypothesize that these NORs would also cluster together in 3-D genome structure. To investigate this, we used Hi-C to reveal more information of genome structure. Our findings revealed that the insertion and removal of rDNA arrays did affect genome structure dramatically (Figure 5C and Figure S8). As previously reported (Mercy et al., 2017), the rDNA array splits *chrIII* or *chrXV* into two non-or low-interacting regions, whereas the removal of the native rDNA locus on *chrXIIR* results in increased interactions between its flanking regions (Figure 5C-D and Figure S8). Our 3D representations of contact maps reveal that the conformation of *chrXII*, *chrIII* and *chrXV* exhibited obvious changes in III-XII-XV and III-XV, and that NORs located on different chromosomes exhibited a tendency to cluster together (Figure 5C).

If the two NORs come into closer proximity in the 3-D structure, there will be more interactions between corresponding flanking regions detected. To quantitatively evaluate the changes, we analyzed the interactions between the 50 kb regions surrounding the rDNA insertion sites on *chrIII* and *chrXV* (the *III-XV* contact type, Figure 5D). Following normalization to BY4742, we found that the *III-XV* type interactions in III-XV-XII and III-XV were significantly higher than those in XV-XII (Figure 5D). This suggests that the rDNA array on *chrIII* enhances the *III-XV* type interactions in strains carrying an rDNA array on *chrXV*, regardless of whether the native rDNA array on *chrXII* has been deleted or not. In addition, more *III-XV* type interactions were detected in III-XV than in III, supporting the positive impact of the rDNA array on *chrXV* on the *III-XV* type interactions in strains containing an rDNA array on *chrIII*. Taken together, these results illustrate the contribution of rDNA arrays to the clustering of themselves in 3-D genome structure.

### Changes in genome structure caused by the alteration of rDNA arrays have little influence on global gene transcription

Currently, the relationship between genome structure and its function in gene regulation remains incompletely understood (Oudelaar and Higgs, 2021). To investigate the impact of additional rDNA arrays on gene expression, we conducted a comparative analysis of the transcriptome between the haploid strains carrying additional rDNA arrays and BY4742 (Figure 6, Table S2). Our analysis revealed that only a small number of DEGs were identified in XV-XII, III-XII and III-XV-XII (Figure 6A-C), and there was no shared DEGs in all three strains (Figure 6D). These results indicated that the presence of additional rDNA arrays does not significantly affect gene expression.

**Figure 6.**
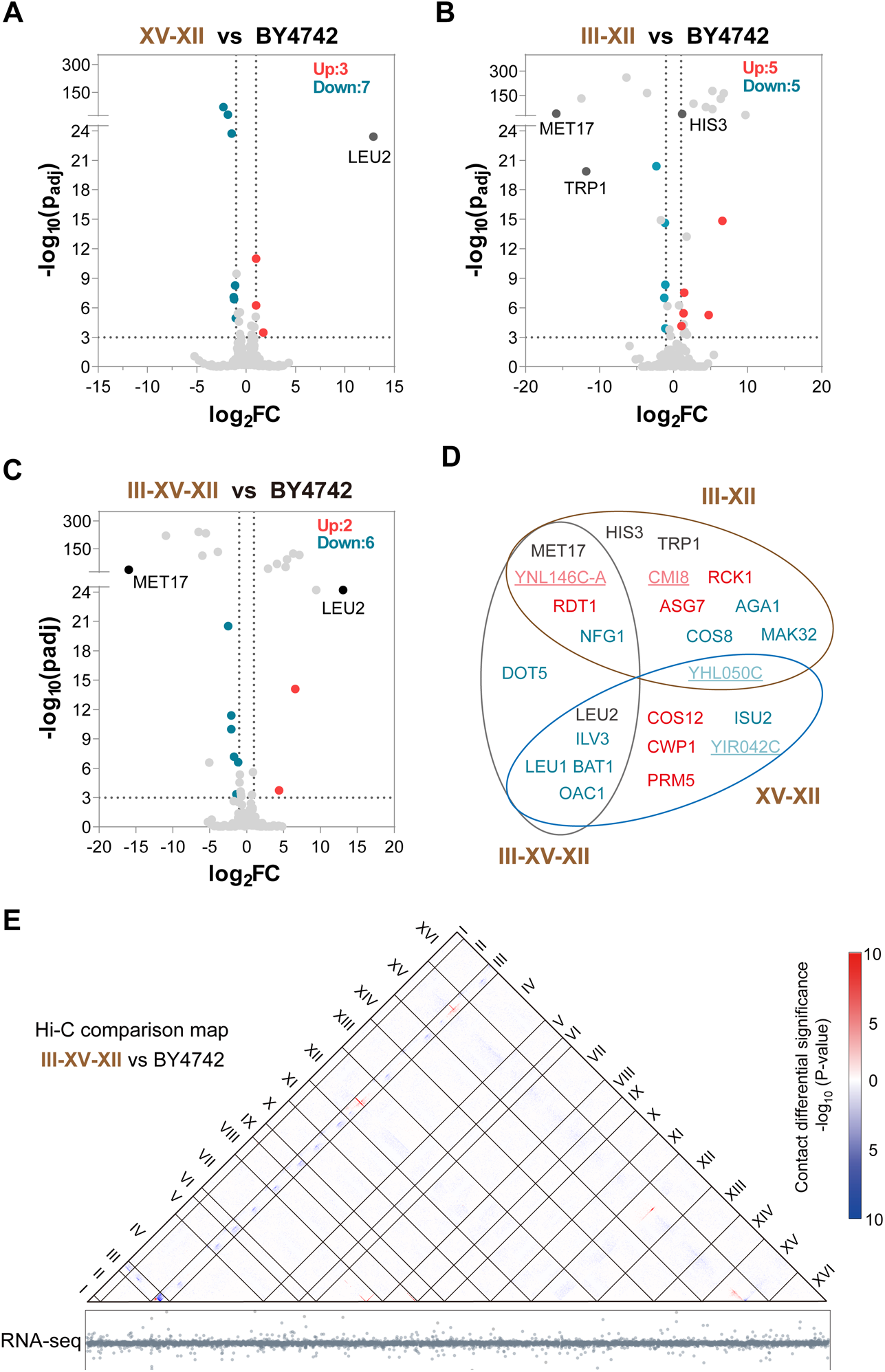
The additional rDNA arrays did not disrupt global gene transcription. (A-D) Transcriptome analysis of XV-XII, III-XII and III-XV-XII. Fold changes (FC) of transcription level were calculated by using BY4742 as wild type control. Black dots: marker genes. Red genes: up-regulated genes. Blue genes: down-regulated genes. Underlined genes: the genes annotated as uncharacterized protein coding genes (SGD, https://www.yeastgenome.org/). (E) The Hi-C comparison map (up) and the RNA-seq results (down) of the strain III-XV-XII and wild type. The location of genes in RNA-seq corresponds to those in Hi-C map. The Hi-C comparison map presents the contact differential significance in corresponding location with -log_10_(p-value). The red colors show the sites with increased contact frequency, and the blue colors present the ones with reduced contact frequency. For the RNA-seq results, the y axis is the log_2_(FC) of expression level.

To better understand the effects of altered genome structure on global transcriptome, we compared the RNA-seq data and Hi-C contact map of III-XV-XII to those of BY4742. The III-XV-XII strain possessed two ectopic rDNA arrays respectively on chromosome III and XV, which underwent dramatic changes in 3-D genome structure, particularly on chromosome III (Figure 5C). Our analysis did not reveal any clustering of expression perturbation near the positions of ectopic rDNA arrays or on any of the two chromosomes (Figure 6E). These findings suggest that changes in genome structure resulting from the alteration of rDNA arrays have little influence on global gene transcription.

## Discussion

In this study, we deployed the CAT method to efficiently construct rDNA arrays with modified sequences at targeted genomic loci. We established another strategy to engineer rDNA copy number using the Cre-loxPsym system. Combining the reported *hyg-1* (Zhang et al., 2017), we successfully induce the increase and decrease of rDNA copy number as we want (Figure 3, S2C and S4). Especially, we reduced the rDNA copy number in a haploid strain to only eight copies. We also successfully constructed haploid strains with double or triple rDNA arrays on different chromosomes. Although the changes of the positions and number of rDNA arrays led to changes in 3-D genome structure, the additional rDNA arrays exerted little influence on global gene expression, growth phenotypes or nucleolar structure. Our results highlight that yeast cells are able to endure dramatic changes to their rDNA organization.

Redundancy is a common feature of genomes. One challenging aim of synthetic genomics is to devise a minimal genome that contains only the genes required to sustain free-living self-replication, to help understand the core functions of life (Vickers, 2016). After several successful endeavors to synthesize and minimize bacterial genomes, the maximum genome reduction of 49% was achieved in mycoplasma cells (Hutchison et al., 2016; Kurasawa et al., 2020; Venter et al., 2022). Due to the larger genome size and gene number, the first project to synthesize a eukaryotic genome, the Sc2.0 project, was initiated in 2006 and is still ongoing (Dai et al., 2020; Richardson et al., 2017). Upon completion of the Sc2.0 project, the native genome will undergo a reduction of approximately 8% in size. The rDNA repeats themselves (100 to 150 copies, ∼9 kb) occupy about 8% to 11% of the entire yeast genome (∼12 Mb). In this study, the rDNA copy number was reduced to just 8 copies (Figure 3). It demonstrates a successful reduction in genome size of approximately 8% achieved solely through the specific removal of rDNA repeats. In addition, the extremely low copy number observed, which approximates those of prokaryotes (typically less than seven copies) (Lavrinienko et al., 2021), suggests potential similarities in fundamental requirements between eukaryotic and prokaryotic life forms. Due to the growth defects and reverse duplication of rDNA region on *chrIII* in S2 and S3, we did not conduct further engineering. Taken the defects resulting from the synthetic array on *chrIII* into consideration, the minimal rDNA copy number for viability may be much lower with a fully functional rDNA array on *chrXII*. And cells without transposons, such as the final Sc2.0 strains, may be useful in constructing more stable low-copy strains for further systematic investigation.

Besides rDNA copy number, we have also modified the number of rDNA arrays. Notably, strain III-XV-XII, containing two extra rDNA arrays, exhibited a substantially shorter doubling time in nutrient-rich media in comparison to BY4742, III-XII and XV-XII (Figure 4C and S6B). However, III-XV-XII, III-XII and XV-XII all showed nearly identical transcriptomes to BY4742 (Figure 6). Furthermore, the specific DEG identified in III-XV-XII, *DOT5* (Figure 6D), has not been previously linked to the regulation of cell growth during the exponential phase. Given the central role of ribosomes in protein translation, the benefits conferred by the two additional rDNA arrays may possibly be manifested predominantly at the protein level, rather than at the transcription level.

As the organelle formed by rDNA arrays, the nucleolus is the most prominent nuclear body with the primary role as the site of ribonucleoprotein particle assembly for ribosome biogenesis (Lafontaine et al., 2021). Previous findings have indicated that the splitting of rDNA arrays to two chromosomes does not affect the formation of a single nucleolus in yeast strains (Hult et al., 2017). These authors have proposed that cross-linking within rDNA, mediated by proteins that bind to multiple sites in rDNA, drives the separation of rDNA from other chromatin, and speculated that irrespective of their positions across the entire genome, multiple rDNA loci would converge to form a single nucleolus. In this study, our strains with two to three NORs all showed a single-nucleolus structure, regardless of the number, position, or length of rDNA arrays (Figure 5). In the 3-D genome structure, NORs situated on distinct chromosomes exhibited a proclivity to cluster together (Figure 5). These findings align with the phase separation hypothesis mentioned above.

In addition, concerted evolution of rDNA is also intriguing due to the highly uniform sequence observed in all repeats within a species, despite interspecific differences in the sequences, regardless of the number of rDNA arrays (Eickbush and Eickbush, 2007). Previous findings suggest that inter-chromosomal genetic exchange and intrachromosomal homogenization may mediate this phenomenon, with the latter occurring at a much higher rate (Eickbush and Eickbush, 2007; Schlotterer and Tautz, 1994; Worton et al., 1988). While existing studies have investigated homogenization dynamics within a single rDNA array in yeast genomes (Eickbush and Eickbush, 2007; Ganley and Kobayashi, 2007, 2011), there remains a paucity of systems to track inter-chromosomal recombination events between different rDNA arrays. In this study, we constructed strains with multiple rDNA arrays containing different sequences and detected only a single instance of a chimeric event in these strains, indicating high homogenization within arrays (Figure 4, S4 and S5). Using experimental evolution, our engineered strains, such as III-XV-XII, will offer an impressive opportunity to track the spatiotemporal dynamic of inter-chromosomal recombination across thousands of generations, a task that is challenging for higher organisms.

## Acknowledgements

This work was supported by grants from National Key Research and Development Program of China (2018YFA0900100), National Natural Science Foundation of China (31725002, 31800069 and 31800082), Shenzhen Science and Technology Program (KQTD20180413181837372), Guangdong Provincial Key Laboratory of Synthetic Genomics (2019B030301006), Guangdong Basic and Applied Basic Research Foundation (2023A1515030285), Bureau of International Cooperation, Chinese Academy of Sciences (172644KYSB20180022) and Shenzhen Outstanding Talents Training Fund. This work was also supported by UK Biotechnology and Biological Sciences Research Council (BBSRC) grants BB/M005690/1, BB/P02114X/1 and BB/W014483/1, Royal Society Newton Advanced Fellowship (NAF\R2\180590) and a Volkswagen Foundation “Life? Initiative” Grant (Ref. 94 771) to YC. This work was also supported by Tip-top Scientific and Technical Innovative Youth Talents of Guangdong Special Support Program (2019TQ05Y876) to YS. We thank Dr. Stefan Hoffmann for proof-reading this manuscript.

## Author contributions

JD, SJ and YC designed and supervised the study. SJ, ZC, CZ, JZ, WY, CS, SZ performed the experiments. SJ, ZC, CZ, YW, WY, SZ, YC, YS carried out the sequencing and analyzed the data. JD, SJ, YC and YM analyzed the data and wrote the manuscript. All authors contributed to review and editing of the manuscript.

## FIGURE LEGENDS

**Figure S1.**
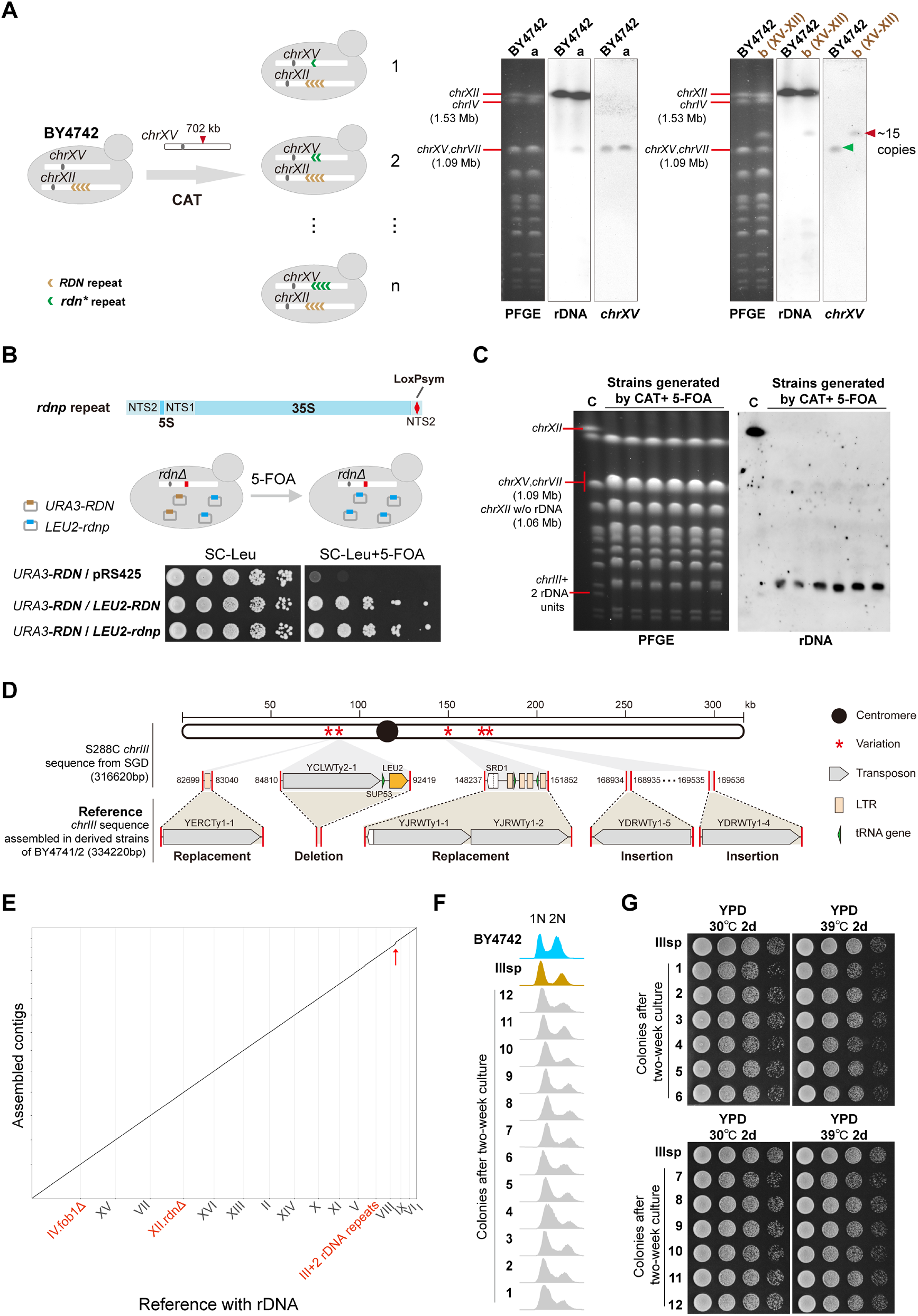
Construction of rDNA arrays through transformation. Related to Figure 1. (A) Construction of an additional rDNA array on *chrXVR* in wild type strain BY4742. Two random strains after CAT, a and b, were analyzed with PFGE and southern blotting. Probes for rDNA, *chrIII* and *chrXII* were used. (B) The plasmid shuffling assay used to test the function of the rDNA repeats with a loxPsym site (*rdnp*). The loxPsym cassette was inserted in the NTS2 region (position -206). (C) PFGE and southern blotting analysis of the random strains generated by CAT and plasmid shuffling. C: the control strain containing two rDNA repeats inserted in *chrIII* and the native rDNA array on *chrXII*. Probes for rDNA were used. (D) The *chrIII* sequences in the derived strains of BY4742 assembled in this study showed several differences to the information from yeast genome database (NC_001135, SGD, https://www.yeastgenome.org) at the positions of Ty-transposons. (E) Dot plot analysis suggested that the synthetic rDNA array has been correctly assembled in *chrIII* of the strain IIIsp. Red arrow indicates the insertion of rDNA repeats. (F) The ploidy of the isolates. Asynchronous log-phase cells were used for analysis. (G) The phenotypes of IIIsp and the twelve isolates after two-week culture.

**Figure S2.**
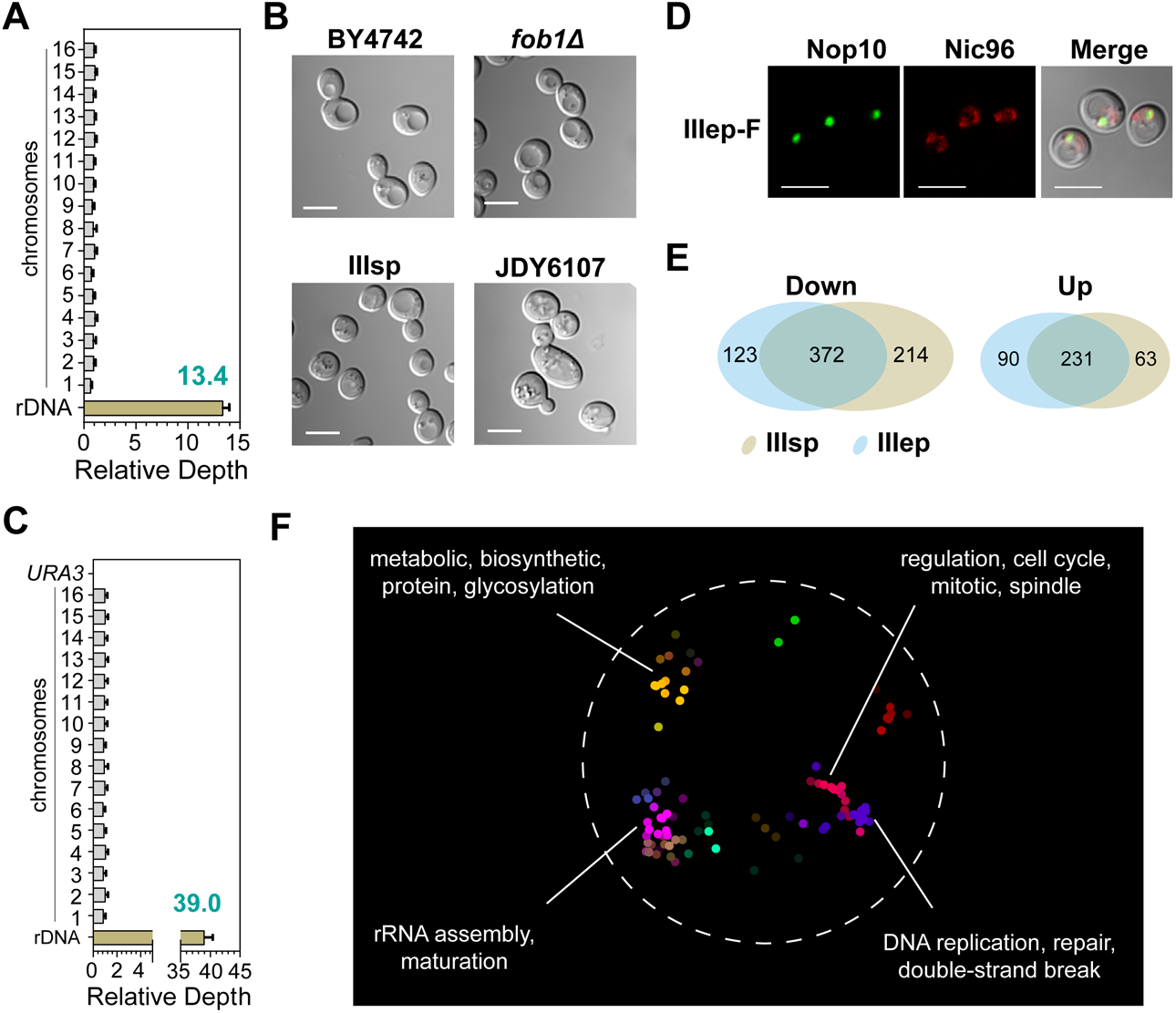
The phenotypes of strains containing a synthetic rDNA array on *chrIII*. Related to Figure 2. (A) The rDNA copy number of the IIIsp-F strain in Figure 2A. Error bars represent SDs. (B) Microscopy pictures of the strains. (C) The rDNA copy number of the IIIep-F strain. The error bar means SD. (D) Microscopy pictures of IIIep-F. IIIep-F showed a normal nucleolar structure. (E) Comparison of the DEGs in IIIsp and IIIep. Down: down-regulated DEGs. Up: up-regulated DEGs. (F) Enrichment landscape of the 603 shared genes (both down-and up-regulated DEGs) in (E) using SAFE (Baryshnikova, 2016). Different colors represent different functional domains. Main domains were labeled with tags.

**Figure S3.**
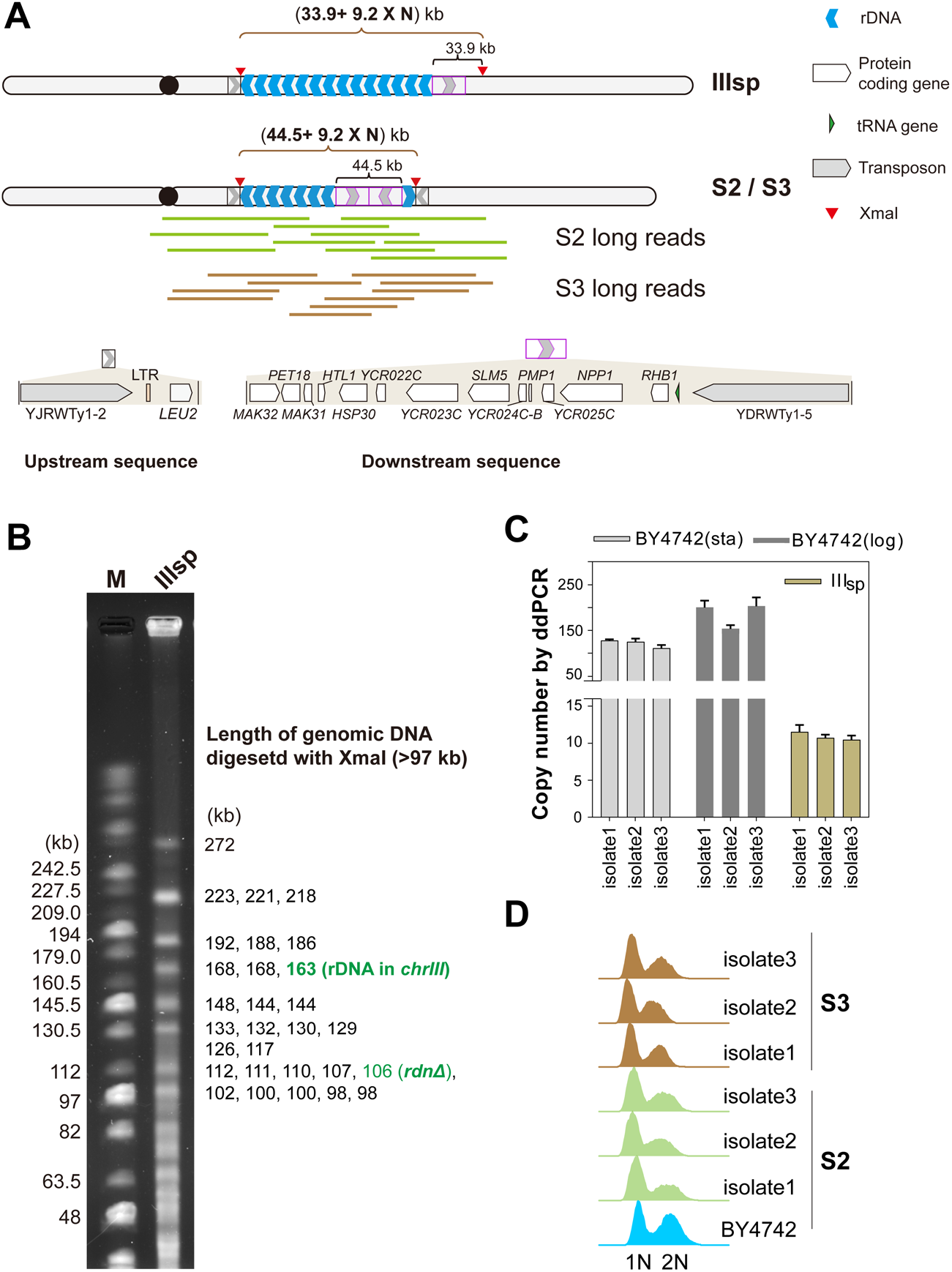
Evaluation of rDNA copy number and genome ploidy. Related to Figure 3. (A) Diagram of *chrIII* assembled in IIIsp and S2/S3 using sequencing data. Red arrowheads: XmaI recognition sites flanking the rDNA region. Reverse duplication of the region including rDNA repeats mediated by Ty transposon was detected in S2 and S3. Representative long reads of S2 and S3 were shown below. Original ID of these reads were listed in Table S4. (B) PFGE results of digested genomic DNA of IIIsp. NEB MidRange PFG Marker (M) was used. The theoretical length of DNA fragments longer than 97 kb is listed on the right. The fragments containing rDNA in *chrIII* or the *rdnΔ* locus in *chrXII* are marked by green color. (C) Assessment of rDNA copy number with ddPCR. Three isolates streaked from each strain were analyzed. Error bars represent the SDs of four technical repeats. Asynchronous BY4742 cells at stationary phase (sta) and logarithmic phase (log) were analyzed. (D) Ploidy of the strains used in Figure3E. The haploid control BY4742 was used.

**Figure S4.**
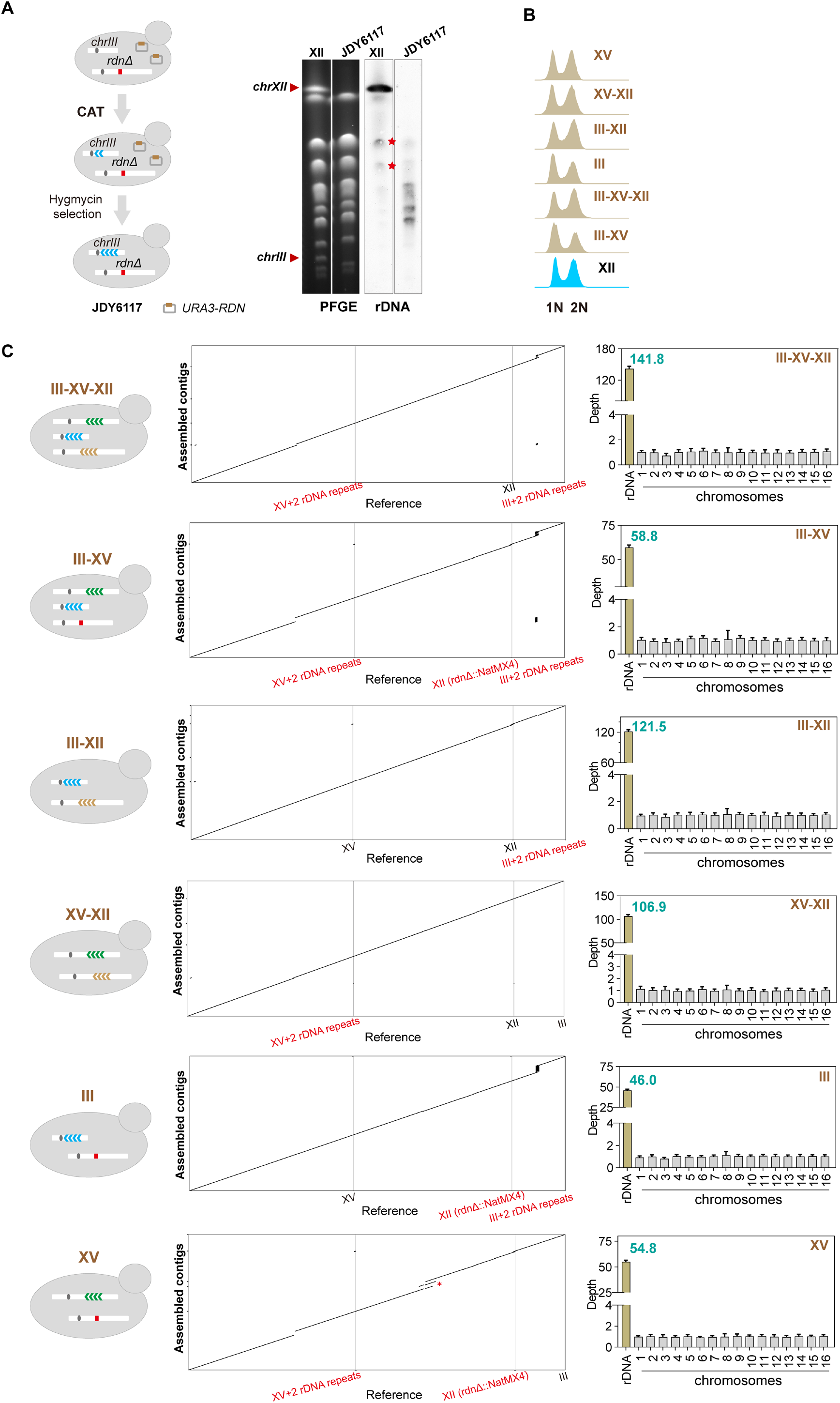
Construction and verification of strains with different number of rDNA arrays. Related to Figure 4. (A) Construction of the strain with an expanded synthetic rDNA array on *chrIIIR* (JDY6117). PFGE and southern blotting analysis of the strain were shown on the right. Probes for rDNA were used for southern blotting. WT: BY4742. Red star: unspecific bands. (B) Genome ploidy of these strains. (C) Verification of strains in Figure 4A by Nanopore sequencing. Dot plot analysis of chromosome III, XV and XII in corresponding strains is shown in the middle. The total rDNA copy number of corresponding strains calculated by sequencing depth is shown on the right. Red star: the sequences of these short contigs had been reviewed manually and no discernible differences were observed in comparison to the reference sequences, except the removal of a Ty-transposon.

**Figure S5.**
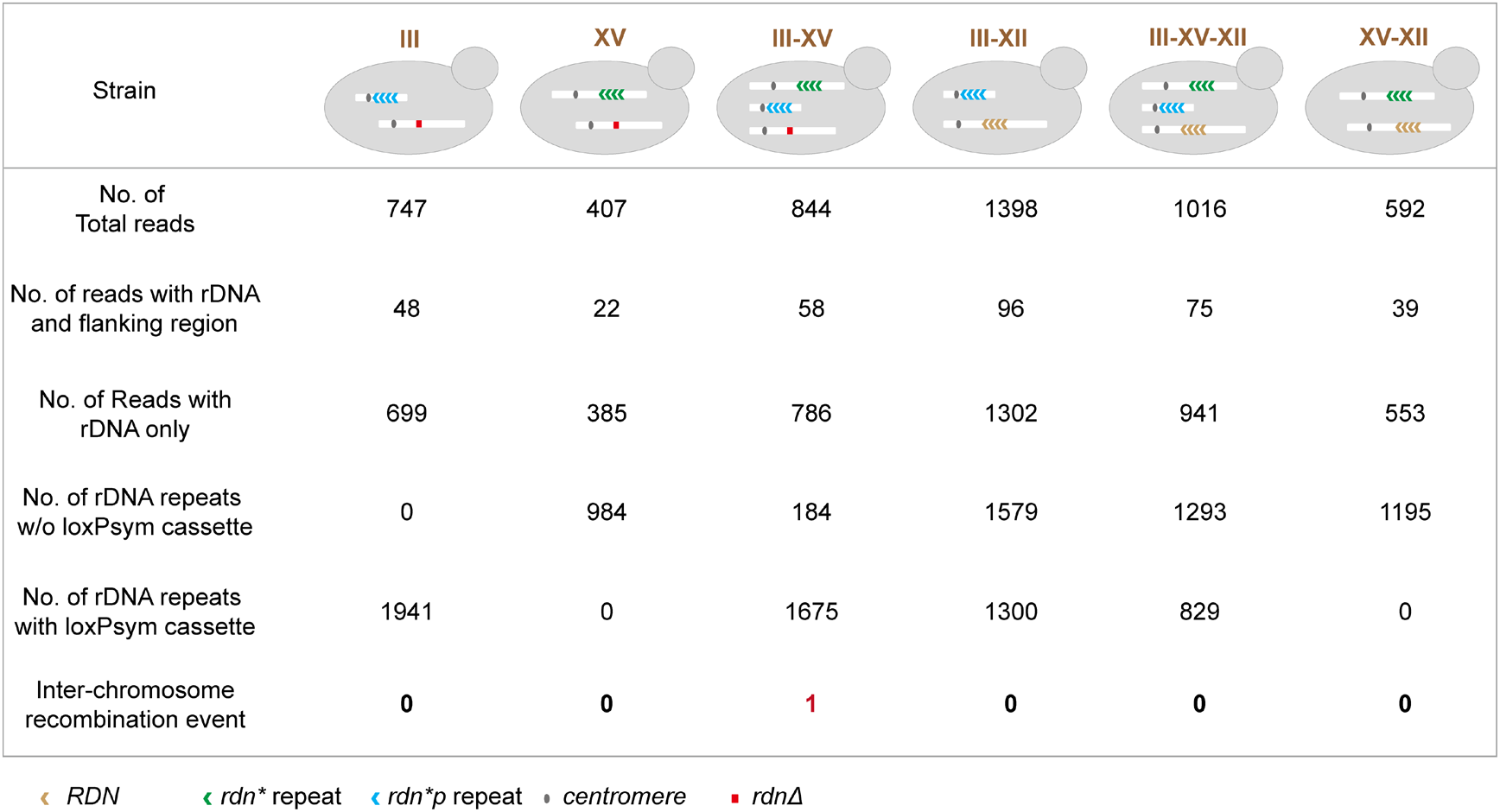
Rare inter-chromosome recombination event between the rDNA arrays. Related to Figure 4 and S4. Total rDNA reads were identified by the alignment of rDNA repeat (∼9kb) in Blast v2.2.26 with the threshold of E-value < 10E-7, identity > 0.65 and coverage > 0.65. The read type and the number of different rDNA repeats were identified by the alignment of the 500bp flanking regions on *chrXII/III/XV*, rDNA repeat (∼9kb) and the loxPsym cassette in Blast v2.2.26 with the same parameters above. Only one chimeric event presents in one rDNA read (one repeat with a loxPsym site vs 4 repeats without loxPsym sites).

**Figure S6.**
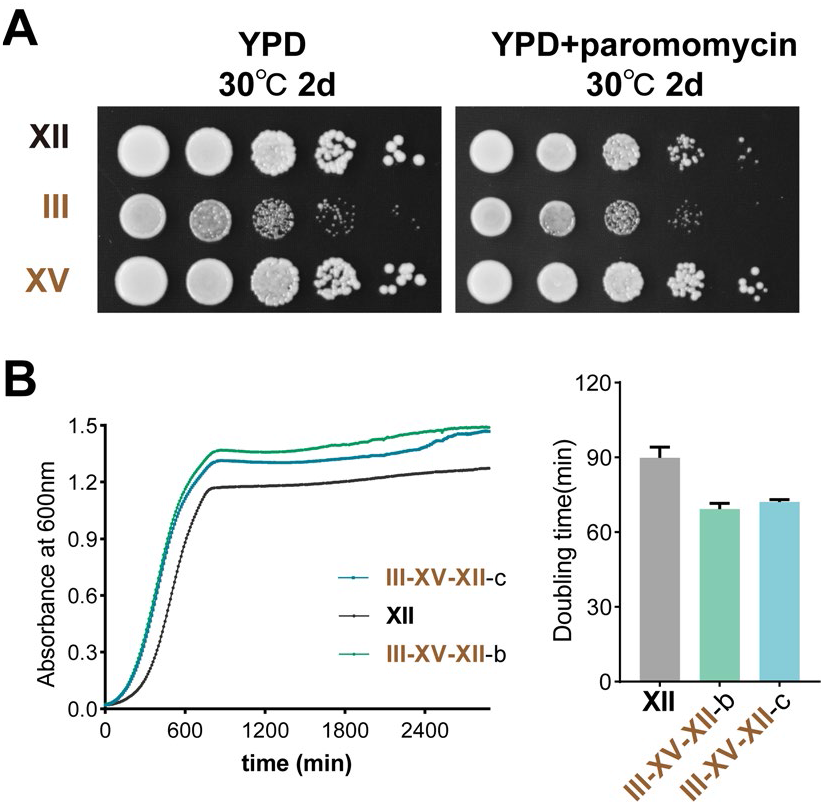
The growth analysis of the strains. Related to Figure 4. (A) The phenotypes of strains were tested on YPD medium with or without paromomycin. Paromomycin: 1mg/ml. (B) Growth of the strains in YPD at 30°C. Two more independent clones for III-XV-XII were analyzed. The mean of three or four technical replicates of each clone was shown for growth curve. The average doubling time is shown in the right. Error bars represent SD.

**Figure S7.**
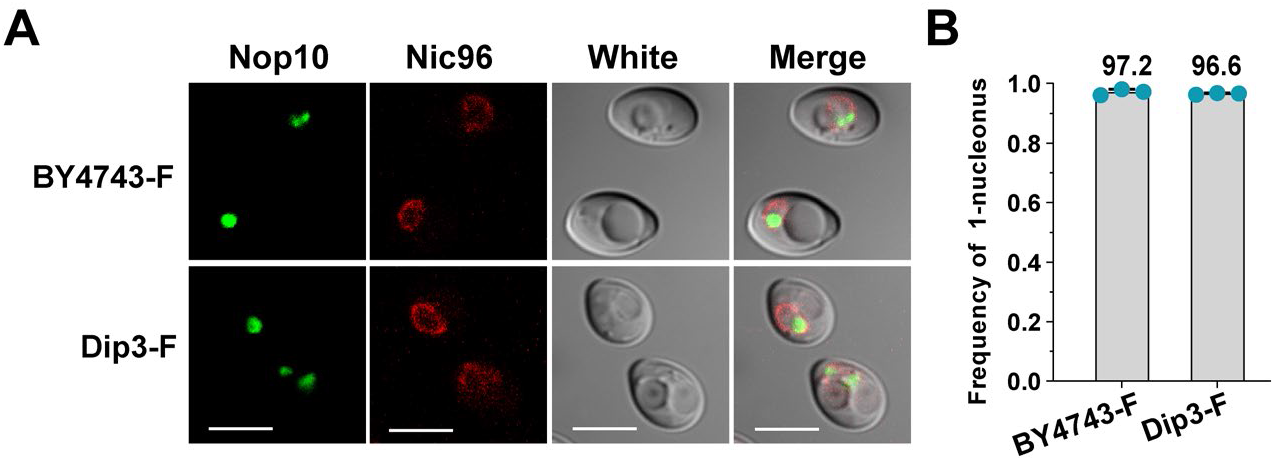
The nucleolus structure of the strains. Related to Figure 5. (A) Microscopy pictures of the strains tagged with Nop10-GFP and Nic96-mCherry. The bar corresponds to 5μm. (B) The frequency of a single nucleolus in different diploid strains. More than 300 un-budding cells were calculated for each clone. Three independent clones were analyzed for each strain (blue dots), with the average frequency being listed above.

**Figure S8.**
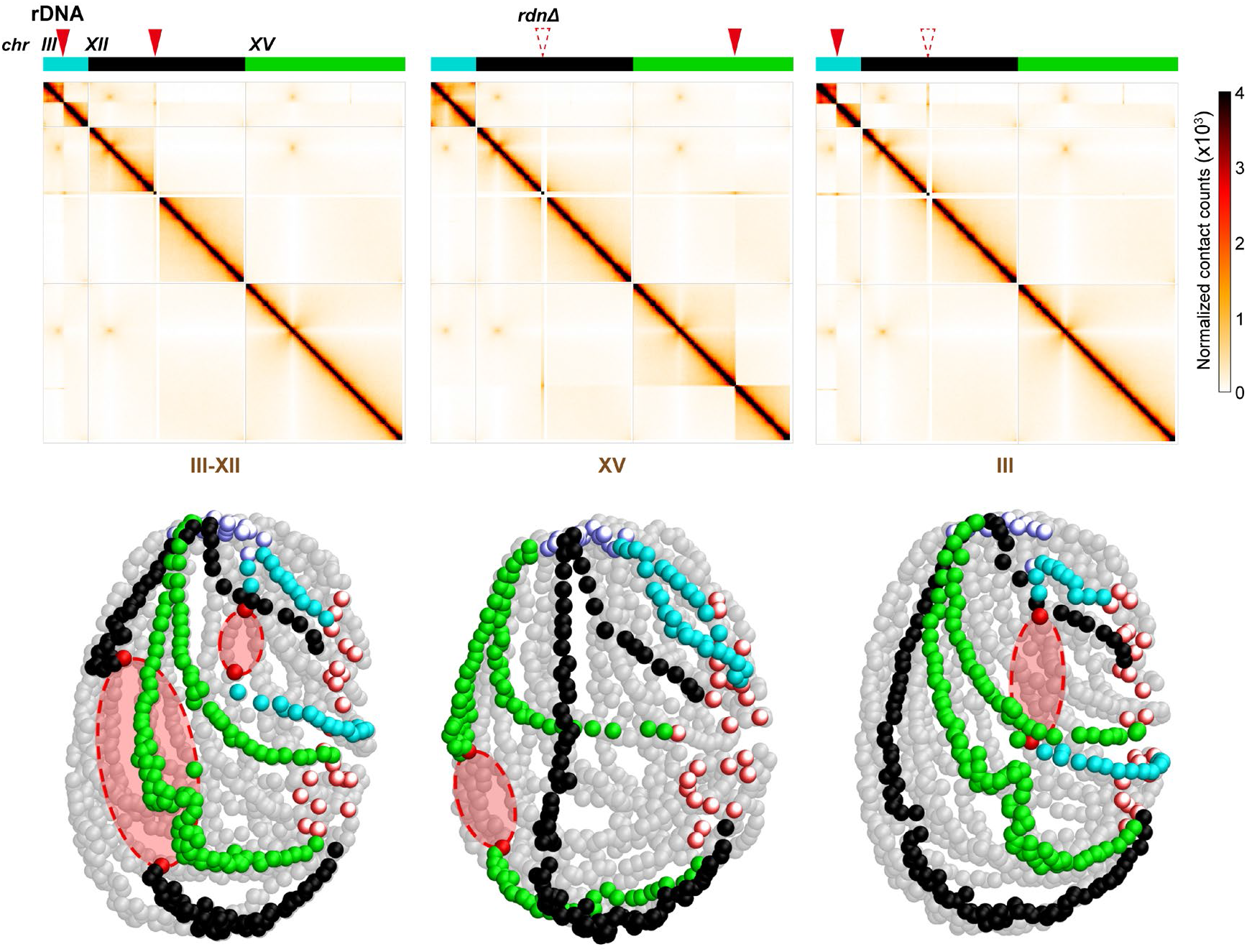
Alteration of the number and position of rDNA arrays affects genome structure. Related to Figure 5. Hi-C contact maps of the three chromosomes (*chrIII*-blue, *chrXII*-black, *chrXV*-green) in corresponding strains and the 3D representations of corresponding contact maps were shown as Figure 5C.

## STAR Methods

### Resource availability

#### Lead contact

Further information and request for reagents and resources should be directed to and will be fulfilled by the lead contact, Junbiao Dai.

#### Materials availability

All the requests for the generated plasmids and strains should be directed to the lead contact and will be made available on request after completion of a Materials Transfer Agreement.

#### Data and code availability

- Nanopore sequencing data, Hi-C data and RNA-seq data have been deposited at SRA with bioproject number PRJNA951483 and are publicly available as of the date of publication.
- This paper does not report original code.
- Any additional information required to reanalyze the data reported in this paper is available from the lead contact upon request.

### Experimental model and subject details

#### Strains and growth media

All strains and plasmids used in this study are listed in Table S3 and key resources table. Oligos used for southern blotting and ddPCR are listed in key resources table. The strain with complete deletion of native rDNA on *chrXIIR,* JDY6106, was constructed and confirmed as before (Zhang et al., 2017). Deletion of *fob1* was mediated by PCR fragments containing the selection marker and flanking sequences as reported (Brachmann et al., 1998). The two fragments of *rdn\** repeats were released from pJD1686 (1 copy) and pJD1687(1.2 copies) through enzymatic digestion (NotI), and co-transformed into yeast cells to construct the rDNA array at the reported locus on *chrXVR* (Zhang et al., 2017). The transformation mixtures were selected on SC-LEU plates. Two candidates after PCR verification of integration were randomly selected for further analysis of the copy number of integrated rDNA repeats.The coordinates for the homologous sequences to the target locus in *chrXV* (NC_001147) : upstream arm in pJD1687,701353-702164; downstream arm in pJD1686, 702169-702955. The two fragments of *rdn*p* repeats were released from pJD1689 (1 copy) and pJD1690 (1.2 copies) through enzymatic digestion (NotI), and co-transformed into yeast cells to generate the rDNA array on *chrIIIR* in this study. For the construction of IIIsp, the transformation mixtures were plated on SC-LEU plates for two days. Colonies on SC-LEU plates were directly replicated to SC-LEU+5-FOA plates to identify the ones without the *URA3-RDN* plasmids. Six colonies on SC-LEU+5-FOA plates were randomly picked for PFGE and southern blotting analysis (Figure S1). The coordinates for the homologous sequences to the target locus in *chrIII* (NC_001135) : upstream arm in pJD1690,151922-152371; downstream arm in pJD1689, 152379-152765. For phenotypic analysis of strains, cells were cultured in YPD medium or YPD medium containing 1mg/ml paromomycin.

### Method details

#### Spot assay

Single colonies were cultured overnight and then subcultured into fresh medium (start OD_600_=0.1) for about another 6 h. Cells with the same optical density at 600 nm (OD_600_) were serially diluted by 10-fold. Then cells were spotted onto YPD plates and incubated at 30°C or 39°C. YPD medium containing 1 mg/ml paromomycin were used.

#### Pulsed-Field Gel Electrophoresis and Southern blotting

Cells from single colonies were cultured in YPD for about 3 days at 30°C to reach stationary phase. ∼2×10^8^ cells were harvested and resuspended in 200µl 0.6% low melting-point agarose and 8 µl 5 mg/ml zymolyase 100T solution (Amsbio, 120493-1). The mixtures were transferred to BioRad plug molds to prepare plugs. The solidified plugs were incubated in 400 µl EDTA solution (500 mM EDTA, 10 mM Tris, pH 7.5) at 37°C overnight. Then 80 µl of 2.5 mg/ml Proteinase K solution (5% sarcosyl, 2.5 mg/ml proteinase K in 500 mM EDTA pH 7.5) was added and incubated at 50°C for at least 5 h. Then the plugs were washed with TE buffer (2 mM Tris, 1 mM EDTA, pH 8.0) more than three times. For the full-length chromosomes, plug samples were resolved on a 0.9% agarose gel with 0.5x TBE for 22-24 h at 14°C on a BioRad CHEF Mapper XA Pulsed Field Electrophoresis System. The voltage was 5.5 V/cm, at an angle of 120° and switch time from initial 50s to final 2 min. For the digested chromosomes, agar plugs were first washed with 500 µl 1x CutSmart buffer and then equilibrated with 800 µl the same buffer for 24 h at 4°C. XmaI digestion was performed in a final volume of about 500 µl: one plug, 50 µl 1x CutSmart buffer, 10 µl XmaI (100U, NEB, R0180L) and ddH_2_O. The tubes were incubated at 4°C for 8 h, followed by an additional incubation at 37°C for 16 h. Cold TE buffer was used to stop the reaction and to wash the plugs three times. Then plugs were resolved on a 1% agarose gel in 0.5x TBE for 24 h at 14°C, with a voltage of 6 V/cm at an angle of 120°, and a switch time from initial 1s to final 20s.

After pulsed-field gel electrophoresis, DNA in the agarose gel was transferred to Hybond-N^+^ membrane (GE Amersham, RPN303B) and UV crosslinked. The preparation of probes labeled with digoxigenin (DIG) and hybridization were conducted using DIG-High Prime DNA Labeling and Detection Starter Kit II (Roche). After hybridization, the membrane was exposed at 477 nm. The probes before labeling were prepared by PCR using the primers listed in key resources table .

#### Doubling Time Assay

The measurements of growth curve were performed as reported previously (Hu et al., 2022). Briefly, the log-phase cells were diluted with fresh medium to the same density (OD_600_=0.05). 100 µl of diluted cells were added to Costar clear polystyrene 96-well plates. Three technical replicates were analyzed for each strain. The 96-well plate sealed with Breathe-Easy membrane (Sigma, MKBZ0331) was cultivated in an Epoch2 microplate photometer (BioTek) at 30°C for 36 h in YPD. The OD_600_ of each well was recorded every 10 min. Doubling times were calculated using GraphPad Prism 8.

#### Genomic DNA preparation, sequencing and analysis

Cells from single colonies were cultured into 5 ml YPD medium. 1 ml overnight culture was diluted into 50 ml fresh YPD and cultured for another about 60 h to reach stationary phase. ∼3×10^9^ cells were collected and washed once in 4 ml cold TE buffer. Cell lysis and DNA extraction were completed using QIAGEN Genomic-tip 100/G (QIAGEN, 19060 and 10243). Then DNA was eluted and precipitated with isopropanol. Finally, DNA pellets were washed twice with 1.2 ml 70% EtOH and dissolved in 100 µl TE buffer. DNA was quantified using Qubit. Sequencing library was prepared with Ligation Sequencing Kit and Flow Cell Prime Kit (Oxford Nanopore Technologies, SQK-LSK109 and EXP-FLP002).

ONT software MinKNOW was used to generate FastQ files. Short (read length < 500) and low-quality (average read quality score < 7) reads in FastQ files were filtered by NanoFilt (De Coster et al., 2018). NGMLR was used to map reads to the reference genome (Sedlazeck et al., 2018) and Canu 2.1.1 was used to assemble reads (∼70 x) into contigs (Koren et al., 2017). Mosdepth was used to generate coverage data (Pedersen and Quinlan, 2018). Copy number of rDNA was calculated by dividing the rDNA coverage by the average coverage of the yeast wild-type genome (rDNA region removed). Mummer-nucmer 4 was used to align contigs to reference sequences with default parameters (Kurtz et al., 2004), generating the data to create a collinearity dot plot by R script.

#### Puromycin incorporation assay and western blot

Puromycin incorporation assay was performed as before (Hu et al., 2022). Briefly, the log-phase cells were diluted to the same density (OD_600_=0.5). Then puromycin was added to the cells with a final concentration of 2 mM and incubated at 30°C for 1 hour. 2 OD cells were collected and lysed using alkali treatment and boil method (Kushnirov, 2000). Equal volumes of protein samples were separated on 10% sodium dodecyl sulfate polyacrylamide electrophoresis gels and transferred to 0.22 µm PVDF membranes for detection. The anti-puromycin monoclonal antibody (Merck, MABE343) was used for detection of puromycin-incorporated proteins. A β-actin (PTM BIO, PTM-5018) was used as the loading control.

#### RNA extraction and electrophoresis assay

Single colonies were cultured overnight and then subcultured into fresh medium (start OD_600_=0.1) for another about 6 h. Then 15 OD of cells were harvested and washed twice with pre-cooled PBS. After removal of liquid, cells were frozen with liquid nitrogen and stored at -80°C before extraction. Cell pellets were gently suspended in 600μl TRI Reagent (Sigma, 93289) and 200μl glass beads were added, followed by cell disruption with a beads beater for 3 min. Then the liquid mixture was collected and extracted with chloroform. After centrifugation at 12,400 rpm at 4°C for 10min, the upper layer was collected and precipitated with isopropanol. The pellets were washed once with cold 75% ethanol and dissolved in 100μl DEPC-treated H_2_O. RNA with the same volume was incubated with DNase I for 15 min at 37°C. Then total RNA was recovered using a Monarch RNA Cleanup Kit (NEB, T2040L). Purified RNA was heated with formamide at 65°C for 5 min, electrophoresed using 1.2% agarose gel and subsequently stained with SYBR safe.

#### Transcriptome analysis

RNA sequencing libraries were prepared and sequenced by commercial company (Beijing Novogene Bioinformatics Technology Co., Ltd.) using Next® UltraTM RNA Library Prep Kit for Illumina®. Raw data in fastq format were processed through in-house perl scripts. In this step, clean data were obtained by removing the reads with adapter sequence, ploy-N or low-quality. HISAT2 was used for reads mapping to the reference genome from genome website (http://ftp.ensembl.org/pub/release-99/fasta/saccharomyces_cerevisiae/) directly. FeatureCounts v1.5.0-p3 was used to count the reads mapped to each gene. Differential expression analysis was performed using the DESeq2 R package. The resulting P-values were adjusted using the Benjamini and Hochberg’s approach for controlling the false discovery rate. Differential expression in this paper was defined as |log2 (Fold change) | > 1 and -log10 (Adjusted p-value) >3. False positive results were inferred by the following rules and removed: dubious genes, marker genes and mating type related genes (Table S2).

#### Microscope imaging for nucleolar morphology

Single colonies were cultured in YPD overnight and subcultured into fresh medium (start OD_600_=0.1) for another 5 hours. Cells were washed twice with ddH_2_O. Then the cells were resuspended in ddH_2_O and dropped on slides for imaging with a Nikon A1 confocal microscope under a 60× objective.

#### Flow Cytometry analysis

Flow cytometry analysis of yeast cells was performed as before (Hu et al., 2022). Briefly, asynchronous log-phase cells were collected and fixed with 70% ethanol for 1 hour at room temperature and resuspended in 50 mM sodium citrate (pH 7.0). After sonication on ice, samples were resuspended in 50 mM sodium citrate (pH 7.0). Then cells were treated with RNaseA (0.25 mg/ml) for 3 hours at 50°C and washed once with 50 mM sodium citrate (pH 7.0). And propidium iodide (16 µg/ml) was added to the cells and incubated at room temperature for 30 min. Samples were analyzed with BD FACSCelesta flow cytometer.

#### Induce Cre-mediated recombination

The cells of IIIsp were transformed with the Cre plasmid pJD1691 or the control plasmid pRS413. The transformation mixtures were directly cultured in SC-HIS liquid medium at 30°C for two days without plating. The cultures were diluted (1:100) and cultured in fresh SC-HIS at 30°C for two more days. Then the cells were diluted to OD_600_=0.1 with SC-HIS+2%Raffinose /0.1% Glucose medium and cultured for 4 hours at 30°C. Then, a proper volume of 20% galactose was added to get the final concentration of 2%. After induction at 30°C for 2 hours, cells with the same OD_600_ were serially diluted by 10-fold. On one hand, the diluted cells were spotted on SC-HIS and YPD plates. Colonies of the Cre-plasmid group showed obvious variations of colony size on both plates, while the control group did not. On the other hand, the diluted cells of the Cre-plasmid group were plated onto YPD plates to get about 200 single colonies per plate. After replication to SC-HIS and YPD plates, we got many *his-* colonies. 48 *his-* colonies were randomly picked for growth analysis using the spot assay on YPD. Six *his-* clones which grew either faster or slower than IIIsp on YPD were randomly chosen for further analysis.

#### Measurement of rDNA copy number by ddPCR

Puromycin incorporation assay was performed as before (Hu et al., 2022) with minor modifications. Yeast cells were collected, washed with 0.5 ml ddH_2_O and resuspended into 100 µl yeast breaking buffer (10 mM Tris-Cl, pH 8.0, 100 mM NaCl, 1 mM EDTA, pH 8.0, 2% (v/v) Triton X-100, 1% (w/v) SDS). 100 µl of 0.5 mm glass beads (Biospec, 11079105) and 200 µl of phenol/chloroform/isopropanol (25:24:1) were added into the 1.5 ml tubes with cells and vortexed at 2000 rpm for 10 min. 100 µl ddH_2_O were added into the tube. After mixing thoroughly, the tubes were centrifuged at 14680 rpm for 10 min. The supernatant was transferred into a new 1.5 ml tube containing 500 µl 100% ethanol. Then the tubes were chilled at -20°C for 15 min before centrifugation (13000 rpm, 5 min, 4°C). The pellet was washed with 500 µl 75% ethanol and dried via a vacuum pump (Eppendorf AG 22331 Hamburg, 45°C, 3 min). Genomic DNA was dissolved in 100 µl ddH_2_O. 90 µl of DNA were treated with 1 µl restriction endonuclease EcoRI-HF (New England Biolabs, R3101S) and 0.4 µl 10 mg/ml RNase A for 1 hour at 37°C, then purified via PCR Purification Kit into 30 µl.

For ddPCR, the genomic DNA was diluted to 0.08 ng/µl and 1 µl genomic DNA was used for per 25 µl reaction. Probes and primers used were listed in key resources table. ddPCR was performed according to the manufacturer’s instructions (Bio-Rad QX200). In brief, the mixtures including genomic DNA, probes and primers for 25S rDNA and *TUB1* were prepared and aliquoted into DG8 cartridge (Bio-Rad, 1864008). After transferring the oil droplets into the Droplet Digital^TM^ PCR 96-well plate (Bio-Rad, 12001925), the plates were sealed with pierceable foil heat seal (Bio-Rad, 1814040). Thermal cycling conditions were 95°C 10 min, followed by 40 cycles of 94°C 30 s, 60°C 1 min, and a final 98°C 10 min enzyme deactivation step. Droplets were cycled to the end point and subsequently read using a QX200 droplet reader. Quantification was performed using Quantasoft software. rDNA copy number per haploid genome= (Number of copies of rDNA)/(Number of copies of *TUB1*) (Salim et al., 2017).

#### Hi-C analysis

Hi-C libraries were prepared according to previous studies (Belton and Dekker, 2015). Briefly, 2 x 10^8^ asynchronous log-phase yeast cells were cross-linked for 20 min with 3% formaldehyde at room temperature and quenched with glycine (final concentration: 0.375 M) for 5 min. The cross-linked cells were homogenized by grinding to a fine powder in liquid nitrogen to lyse the cell wall. Endogenous nucleases were inactivated with 0.1% SDS, then chromatin DNA was digested by 100U MboI (NEB, R0147L), labeled with biotin-14-dCTP (Invitrogen, 19518018), and then ligated by 50U T4 DNA ligase (NEB, M0202V). After reversing cross-links, the ligated DNA was extracted using QIAamp DNA

Mini Kit (Qiagen, 51304). Purified DNA was sheared to 300-to 500-bp fragments and was blunt-end repaired, A-tailed and adaptor-added, followed by purification through biotin-streptavidin–mediated pull-down and PCR amplification. Finally, the Hi-C libraries were sequenced on the MGISEQ-2000 sequencing platform (BGI, China).

The raw and normalized contact count matrices of the Hi-C data were calculated at 10 kb resolution by HiC-Pro v3.1.0 (Servant et al., 2015) with default parameters. To detect the contacts between the rDNA repeats and other chromosomes, we retrieved the reads mapped on the rDNA region for the contact count matrices. The normalized contact maps were visualized by pheatmap in R package. The significant changes in contact count matrices were measured by ACCOST (Cook et al., 2020) in the raw contact counts. The 3D genome architecture was constructed using ShRec3D+ (Li et al., 2018) in the normalized chromosomal contacts, and visualized them by PyMoL v2.6.0a0 (Schrödinger, LLC.).

#### Quantification and statistical analysis

Error bars represent SDs. Two-tailed t-tests or nested one-way ANOVA were used to compare different groups in this paper. Differences were considered as statistically significant at p-value<0.05. * P<0.05, ** P<0.01, *** P<0.001, **** P<0.0001.

